# Noninvasive, automated and reliable detection of spreading depolarizations in severe traumatic brain injury using scalp EEG

**DOI:** 10.1101/2022.10.07.511376

**Authors:** Alireza Chamanzar, Jonathan Elmer, Lori Shutter, Jed Hartings, Pulkit Grover

## Abstract

**Background:** Noninvasive detection of spreading depolarizations (SD), as a potentially treatable mechanism of worsening brain injuries after traumatic brain injuries (TBI), has remained elusive. Current methods to detect SDs are based on intracranial recording, an invasive method with limited spatial coverage. Less invasive methods to diagnose SD are needed to improve generalizability and application of this emerging science and to guide worsening brain injury treatments. Here, we demonstrate, for the first time, a signal processing paradigm that can enable automated detection of SDs using noninvasive electroencephalography (EEG).

**Methods:** Building on our previously developed WAVEFRONT algorithm, we have designed a novel automated SD detection method. This algorithm, with learnable parameters and improved velocity estimation, extracts and tracks propagating power depressions, as well as near-DC shifts using low-density EEG. This modified WAVEFRONT is robust to the amplitude outliers and non-propagating depressions on the scalp. We show the feasibility of detecting SD events (700 total SDs) in continuous, low-density scalp EEG recording (95±42.2 hours with 19 electrodes) acquired from 12 severe TBI patients who underwent decompressive hemicraniectomy (DHC) and intracranial EEG that could be used as a ground truth for event detection. We quantify the performance of WAVEFRONT in terms of SD detection accuracy, including true positive rate (TPR) and false positive rate (FPR), as well as the accuracy of estimating the frequency of SDs.

**Results:** WAVEFRONT achieves the best average validation accuracy of 74% TPR (with 95% confidence interval of 70.8%-76.7%), with less than 1.5% FPR using Delta band EEG. Preliminary evidence suggests that WAVEFRONT can achieve a very good performance (regression with R^2^ ≃0.71) in the estimation of SD frequencies.

**Conclusions:** We demonstrate feasibility and quantify the performance of noninvasive SD detection after severe TBI using an automated algorithm. WAVEFRONT can potentially be used for diagnosis and monitoring of worsening brain injuries to guide treatments by providing a measure of SD frequency. Extension of these results to patients with intact skulls requires further study.

## I. INTRODUCTION

This paper is aimed at addressing this question of whether spreading depolarization (SD) waves in the brain can be detected noninvasively. This is a long-standing question in the field of neurocritical care, with limited and contrasting reported results on the feasibility of SD detection using scalp electroencephalography (EEG). An SD is a wave of neurochemical changes in the brain, which propagates slowly (1-8 mm/min) across the cortical surface and suppresses neural activity [1–3]. SDs are caused by a breakdown in the ionic homeostasis across neuronal membranes [4]. Increasing evidence shows that SDs are associated with poor clinical outcomes in traumatic brain injuries (TBIs), stroke, and hemorrhages [4–9]. Experts believe that this association is causal, such that the neurophysiological sequelae of SDs directly worsen secondary brain injury [9–15]. Recent studies have explored the potential of SD as a therapeutic target in subarachnoid hemorrhage (SAH) [9, 16–18] and TBI [19, 20]. Each year, around 2.5 million patients suffer TBI happen in the United States^1^. Recent data on five-year outcomes for patients with moderate and severe TBI in the United States indicate that more than half of these patients experience neurological worsening or death [21]. Therefore, detection of SDs, as a reliable biomarker and a potential therapeutic target, may help improve clinical outcomes.

Previous attempts to detect SDs using EEG have had limited success. In [22], Hartings *et al*. reported amplitude depressions associated with 81% of the intracranially detected SDs, through visual inspection of scalp EEG recordings in severe traumatic brain injury (TBI) patients. In an earlier work [23], Drenckhahn *et al*. reported similar positive results on malignant stroke patients with SAH. In another recent work, some SD-type patterns were observed in scalp EEG recordings of epileptic patients [24], but these are without invasive recordings to serve as a ground truth. By contrast, in [25], Hofmeijer *et al*. monitored 18 stroke and 18 TBI patients using 21-electrode EEG systems, where visual inspection did not reveal any SDs. In [25], patients did not receive craniotomy and there was no invasive recording. Therefore, it is not clear which patients had any SDs. In a commentary paper [26] on this work, Hartings *et al*. provided potential reasons for these reported negative results, including lack of criteria for monitoring of SDs in scalp EEG, spatial low-pass filtering effect of the intact skull in these patients, and using short 1-hour time intervals for visual inspection of the EEG signals in [25]. Using highly compressed EEG recordings in 11-hour time intervals, the authors in [26] reported visual identification of some patterns of SD depressions in a TBI patient, but without any invasive ground truth. This work was followed recently by [9], where the authors monitored 15 TBI and 20 aneurysmal SAH (aSAH) patients, with simultaneous invasive and noninvasive recordings, some of whom received craniotomy. No significant association was found between occurrence of SDs and established delayed cerebral ischemia (DCI) biomarkers in continuous scalp EEG. Thus, on balance it remains uncertain whether noninvasive detection of SDs is feasible, let alone be sufficiently reliable for clinical relevance.

In addition, the feasibility of noninvasive SD detection has been explored using simulations [27–29]. In a related work on real data, we demonstrated the feasibility of localizing *non-spreading* depressions of activity (i.e., *neural silences*) in the brain, using EEG recordings [30, 31], what we call “silence localization”. The insights obtained from silence localization [30, 31] informs our approach here.

This work is the first attempt to explore the feasibility and quantify the performance of noninvasive SD detection in severe TBI patients using an automated algorithm applied to real data, an unmet need in the field. We advance our previous algorithm, WAVEFRONT [28], for automated noninvasive detection of SD events in a group of 12 severe TBI patients who underwent decompressive hemicraniectomy (DHC) surgery, followed by continuous monitoring in ICUs using simultaneous scalp EEG (a low-density EEG system with 19 electrodes at 10-20 standard locations) and intracranial ECoG recordings. WAVEFRONT is an *explainable* automated SD detection framework with intuitive steps and *interpretable* detection outputs and results.

In Section III, we show that WAVEFRONT achieved a reliable SD detection performance using the Delta band EEG recording, with ∼74% average true positive rate (TPR) and less than 1.5% false positive rate (FPR) using cross-validation, as illustrated in Fig. 8. The high TPR attained with a low FPR resolves the *feasibility* question: noninvasive SD detection is possible, at least for patients who have received DHC. However, is this performance sufficient for clinical goals?

For two patients in our dataset, who have no ground truth SD events, the average FPR is 1.7%. This might seem small (and is comparable to the FPR on other patients). However, in raw numbers, this is still associated with a substantial number of false positive events (each event here is a time window in which an SD presence/absence is to be detected). To understand the clinical implications of this and answer the earlier question on the sufficiency of WAVEFRONT’s performance for clinical goals, we perform an additional analysis: prediction of the number of SDs from total minutes of detected SD events. This is inspired by an estimation of how frequently SDs occur in a recent work of Jewell *et al*. [10], where, unlike here, the aim was to automate detection of SDs for *invasive* detection (more details on this work in the next paragraph). Preliminary results included here, albeit with limited data, suggest that WAVEFRONT can reliably estimate the number of occurrences of SDs in long time intervals of 30-hour, with R^2^ ≃0.71 using a regression analysis, as it is shown in Fig. 13, and explained in detail in Section III-D. Around 60% of severe TBI patients experience SD episodes, with a potentially high rate of occurrence [10, 32]. Increasing evidence shows that SDs are a reliable predictor of outcomes in TBI patients [5, 10, 19, 33]. Therefore, we think that WAVEFRONT’s performance is indicative that noninvasive prognostication of worsening brain injuries using SD detection is possible. However, further studies, with more data, are warranted to understand this.

In [10], towards automating ECoG-based SD detection, Jewell *et al*. developed a technique for real-time SD detection using ECoG signals, by combining features from low-frequency bands (e.g., slow potential changes in 0.005-0.5 *Hz*) and highfrequency bands (e.g., reduction of envelope amplitude in 0.5-45 *Hz*). They reported a detection rate of ∼ 79%, with a 0.9% false alarm rate ^2^, in 18 acute brain injuries patients with DHC.

### Clinical relevance of noninvasive SD detection in patients with DHC

As discussed earlier, this work is focused on severe TBI patients with DHC, and a natural question is: is noninvasive and automated SD detection in patients with removed skull parts clinically relevant? The DHC procedure is a part of standard of care for many severe TBI patients [34] to control an elevated intracranial pressure, extract hematoma, and prevent further damage to the brain tissue [35–37]. It is worth noting that following a DHC, the scalp is sutured back over the brain, even though a piece of the skull is missing. Patients who receive DHC are continuously monitored in ICUs after closing the scalp incision. During this period, scalp EEG-based automated SD detection can provide valuable clinical information pertaining to worsening brain injuries. Intracranial monitoring of SDs can provide more information than the presence or absence of these waves in the brain, such as characteristic patterns of SDs including clustered or persistent depressions of spontaneous neural activities [8]. However, scalp EEG has broader spatial coverage than a locally placed ECoG strip. It also provides better spatial resolution in DHC patients in comparison to the intact-skull EEG recordings [35], at least close to locations where the skull has been removed. Further, while procedural risks (e.g., bleeding, infection, etc.) associated with subdural electrodes are infrequent [32, 38], noninvasive EEG would preclude their possibility entirely. Therefore, noninvasive SD detection in severe TBI patients with DHC can prove clinically valuable for improving outcomes.

Furthermore, validation of WAVEFRONT on DHC patients is a good starting point for noninvasive and automated SD detection because: a) intracranial ground truth for SDs can be obtained by placing ECoG electrodes during the DHC procedure, and b) head layers, including skull, meninges, cerebrospinal fluid (CSF), and scalp, have low-pass filtering or *blurring* effects on the scalp EEG signals. This makes the detection and tracking of narrow SD waves challenging [27, 28]. This is less challenging in DHC patients due to the absence of the very low conductivity skull layer. Nevertheless, the challenge is significant: i) relative to ECoG, the signal is more noisy and spatially low-pass filtered, and ii) as the SD wave propagates into the sulcus, its representation in the scalp EEG signals reduces significantly. This breaks the waves, as measurable by EEG, into disconnected components, which we call “wavefronts” [28]. Complex patterns of SDs (e.g., single gyrus [39, 40], semi-planar [40–42], etc.) can make the noninvasive detection of these waves even more challenging.

WAVEFRONT addresses some of the difficulties in noninvasive detection of SD waves in EEG. It breaks down the challenging task of detecting the whole propagating SD wave in the brain using noisy and blurry filtered scalp EEG signals into simpler tasks of detection and classification of disjoint SD wavefronts, following these steps: Power envelopes of the scalp EEG signals at each electrode are extracted, and depressions (power reductions) are detected. These detected depressions are then projected onto a 2D plane to obtain depression wavefronts. Propagating SD wavefronts are then detected and tracked based on their speed and direction of propagation. To estimate the speed and direction of propagation of depression wavefronts on these 2D planes, WAVEFRONT uses a computer vision technique, called optical flow [43, 44]. It then stitches together the detection of these wavefronts over time and space to detect and track SD waves in the brain. This overcomes the challenge related to the effects of sulci and gyri discussed above and enables the detection and tracking of complex patterns of propagation.

Although the simulation results of automated SD detection in [28] are promising, we recognize that WAVEFRONT suffers from certain technical shortcomings and cannot be directly applied to real scalp EEG signals for SD detection: (i) it uses a fixed set of parameters (e.g., depression level/depth threshold, temporal score threshold, and spatio-temporal neighborhood radius), (ii) it is highly sensitive to the amplitude outliers, (iii) it makes an implicit assumption that the power level of normal background brain activity (DC offset of the power envelope) is stable and not changing over time (see Fig. 9 and 10 in [28] for more details), which limits the ability of WAVEFRONT in the detection of depressions, as well as near-DC shifts during propagating SDs in the real EEG recordings, (iv) it does not address the challenges of using a low-density EEG grid, including the high rate of false alarms due to the non-propagating depressions on the scalp that we observe here (see Section II-D), and (v) it estimates the optical flows in pixels on the 2D images, rather than in terms of the physical distances on the scalp, which can introduce errors in estimation of the speed and direction of propagation of SD wavefronts. In this work, we address these limitations of WAVEFRONT by making necessary modifications and improvements (see Section II for details), including designing a training and validation framework for the algorithm to learn an optimal set of parameters through a cross-validation analysis. Other modifications include: (a) designing a rigorous and automated preprocessing pipeline for outlier rejection and pruning the EEG signals, (b) using a power-envelope extraction method which is less sensitive to large-amplitude artifacts (i.e., outliers), (c) extending the depression extraction method to be able to detect DC shifts in the near-DC components ([0.01-0.1] *Hz*), as well as the power depressions in the higher frequency bands (≥ 0.5 *Hz*), (d) defining an “effective propagation measure” along with a learnable threshold on this measure to reject the false alarms of the non-propagating depressions on the scalp, and (e) mapping the estimated optical flows on the scalp spherical surface.

Our study has important clinical limitations. The temporal annotation of an SD event through visual inspection of the ECoG signals may not accurately reflect the actual onset of each SD wave. The reported performance metrics in this study (e.g., TPR and FPR) depend on the temporal annotations of SDs (see Section III-A). This may have resulted in slight over or underestimation of the actual performance of WAVEFRONT in the detection of SD events. In addition, due to the limited spatial coverage of the strip of ECoG electrodes, some of the SDs may be even missed in the “ground truth”. This is a limitation of any technique that uses a ground truth with limited spatial coverage. Also, due to the small number of patients in this study, overfitting to the available SD events is inevitable (see Section III for more details on this issue), which worsens the validation performance. We expect WAVEFRONT to achieve a better average validation performance by using a larger dataset of TBI patients with a higher density of EEG electrodes. WAVEFRONT might also underdetect clustered SDs (more than two SDs in a time interval of 3 hours or less [10, 19]), since it does not detect SDs individually, a limitation that future algorithms can address.

In Section II, we provide the details on the dataset and SD ground truth. In addition, we provide the details of the EEG preprocessing pipeline and the WAVEFRONT algorithm, along with the modifications and improvements we have made. In Section III, we present the performance of WAVEFRONT in noninvasive SD detection across different EEG frequency bands and evaluate the frequency of SDs in large time windows. Finally, we conclude in Section IV, where we discuss some of the limitations of our study and discuss promising direction for future work.

## II. METHODS

In this section, we first provide details on the dataset used in our study, which includes a group of 12 patients hospitalized after severe acute TBI who underwent DHC and cortical strip ECoG electrode placement. We provide details on the intracranial SD ground truth, and then introduce our automated SD detection method, with an emphasis on the explainability of WAVEFRONT by providing the intuition and visualization of the main steps.

### A. Dataset

The dataset we used was obtained as part of a multicenter clinical study that monitored SDs in TBI patients ^3^. Continuous EEG signals were recorded over a few days (95±42.2 hours on average) following DHC using a DC-coupled EEG amplifier (*CNS Advanced ICU EEG Amplifier from MOBERG ICU Solutions*), with a sampling frequency of 256*Hz*, from 19 electrodes placed at 10-20 standard locations. All of the EEG electrodes were referenced to a common reference electrode.

In addition, during the DHC procedure a strip of six monopolar ECoG electrodes (referenced to the same electrode), with an inter-electrode distance of 1cm, was placed on the hemisphere that underwent the DHC, and continuous ECoG and EEG data were recorded simultaneously using the same amplifier. The recorded ECoG signals were visually assessed by a clinical expert (Dr. J. Hartings) to identify and annotate the SD episodes in the dataset. See Section II-B for more information on the SD temporal annotations.

For EEG preprocessing and artifact removal (see Section II-C for details), we use two additional electrophysiological recordings: (i) an Electrocardiogram (ECG) signal recorded at a 500*Hz* sampling frequency, and (ii) a Plethysmogram (PLETH) signal at a 125*Hz* sampling frequency. These signals were recorded using *IntelliVue Bedside (PHILIPS)* patient monitor.

#### Participant

Data from 12 (9 male and 3 female) severe TBI patients were utilized in our study. Two patients had DHC in the left hemisphere and the remaining 10 patients had DHC in the right hemisphere. Eleven patients experienced subdural hematoma (SDH), and one patient had an epidural hematoma (EDH). Detailed information about these patients is included in Table I. All procedures were approved by the University of Cincinnati Institutional Review Board. A legally authorized representative for each patient provided surrogate consent for participation in the initial research study. For visualization, computed tomography (CT) scans of a TBI patient (patient 6, see Table I) with right DHC are shown in Fig. 1, where the locations of DHC and evacuation of hematoma can be seen as asymmetric dark regions in the white thick layer of skull. In addition, the location of the subdural strip is also shown (Fig. 1 right).

**Table I.**
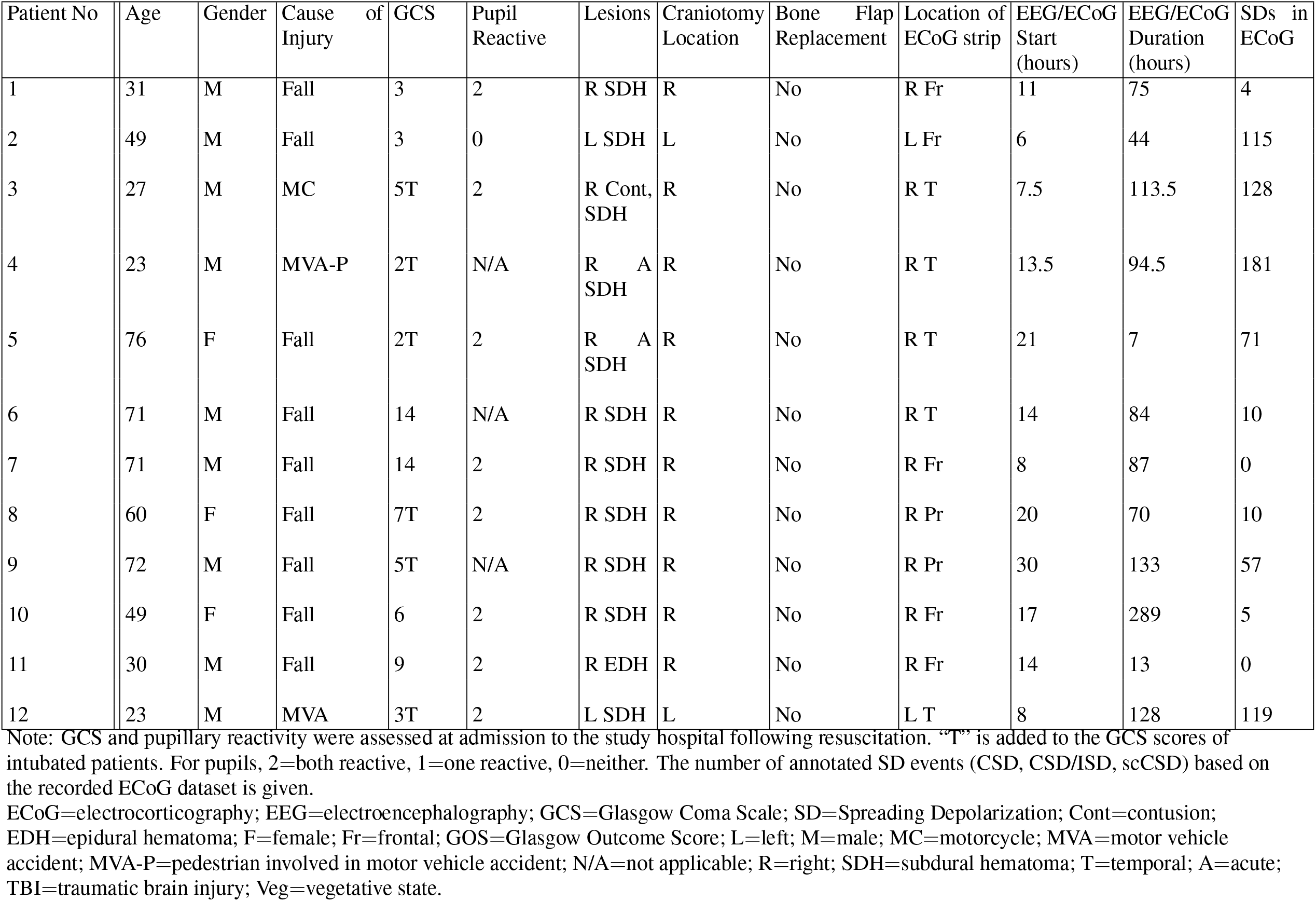
Clinical details and demographic information for 12 TBI patients in the dataset.

**Fig. 1.**
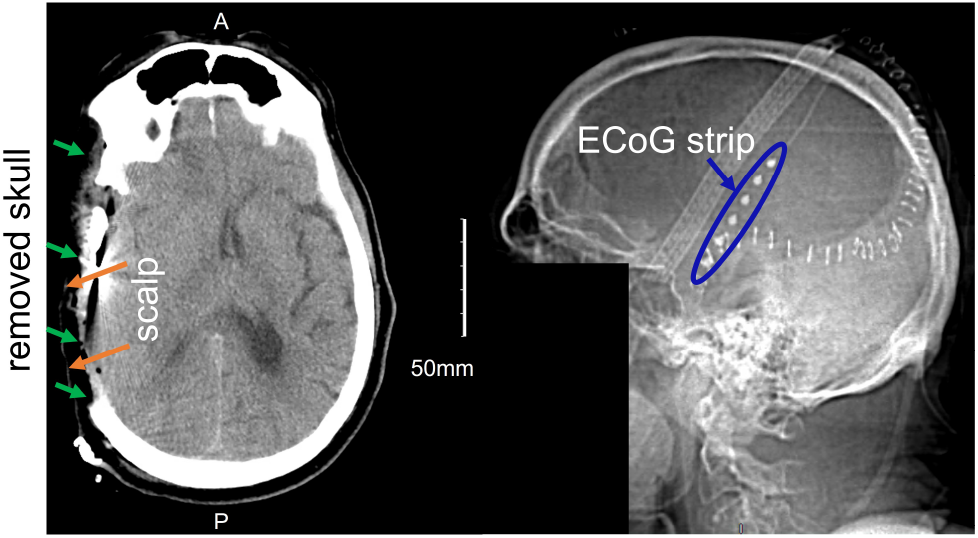
Transverse (left) and right side (right) view of computed tomography (CT) scans of a severe TBI patient (patient 6, see Table I) with right decompressive hemicraniectomy (DHC). The missing portion of the skull, the scalp layer on the DHC site, and the strip of 6 electrocorticography (ECoG) electrodes are also marked. Facial region is stripped away to ensure anonymity of the patient.

### B. SD event temporal annotation based on ECoG signals

Each SD event in these TBI patients was annotated over time by a clinical neuroscientist^4^ through visual assessment of full-band ECoG signals. In this paper, an SD event refers to a *unique* SD wave, as annotated by consideration of, and manifested across, all electrodes of the ECoG strip. Each unique SD wave was annotated at the start of a slow near-DC negative shift (in 0.01-0.1 *Hz*), i.e., slow potential change (SPC), in a chosen ECoG electrode, which is not always the same one even for the same patient. The temporal annotation of an SD event through visual inspection of the ECoG signals may not accurately reflect the actual onset of each SD wave. The reported performance metrics in this study (e.g., TPR and FPR) depend on the temporal annotations of SDs (see Section III-A). Four different types of SD events were annotated: (i) CSD: an event during which there was a cortical spreading depression (CSD) in each electrode that had an SPC, where CSD is a manifestation of spreading depolarization and is defined as a cortical wave of depression in the high-frequency (*>* 0.5*Hz*) ECoG (HF-ECoG) signals [45], (ii) ISD: an event where the HF-ECoG signals at all the participating electrodes with SPC were already flat. These are called isoelectric spreading depolarizations (ISDs) [8], (iii) CSD/ISD: an event where some ECoG electrodes experienced CSD propagation, while other electrodes had ISDs, and (iv) scCSD: an event which was identified as a clear SD in the signal of a single electrode. Although it appeared only on a single electrode, it met the “consensus criteria”, defined by Co-Operative Studies on Brain Injury Depolarizations (COSBID) [8], to be classified as SD. According to COSBID, one of the minimal criteria to score an event as SD is “an event which has a characteristic DC shift associated with spreading depression of spontaneous activity even if DC shift and spreading depression are restricted to a single channel.” The total number of annotated SD events for each patient in the dataset is included in Table I.

In this study, the ground truth SDs are annotated based on a single ECoG strip located in the DHC hemisphere (e.g., see Fig. 1) (no invasive measurements are made in the contralateral hemisphere, i.e., the hemisphere with an intact skull). We observe that the scalp EEG signals from the contralateral hemisphere and ipsilateral hemispheres are significantly different due to the missing part of the skull in the ipsilateral hemisphere (e.g., ipsilateral EEG signals have higher average power, see Fig. 2b). In this paper, we only use ipsilateral scalp EEG electrodes in each patient for SD-related inferences. Excluding the contralateral electrodes is helpful to: (i) tailor the WAVEFRONT algorithm to the SD events that we are certain about (i.e., events that we have a ground truth for) during the training process (see Section II for details), and obtain a more realistic estimation of WAVEFRONT’s performance in noninvasive detection of SDs, and (ii) acknowledge the statistical differences between the ipsilateral and contralateral EEG signals, as ignoring these differences may adversely affect the performance of WAVEFRONT. Because EEG signals tend to be less sensitive to contralateral sources, we expect this restriction to not hurt the performance of our algorithm. Finally, electrodes on the mid-line (Fz, Cz, and Pz) *are* included in our analysis as they can be sensitive to signals on either side. Fig. 2a shows the selected subset of EEG electrodes (11 out of 19 electrodes) for a patient with a right hemisphere DHC.

**Fig. 2.**
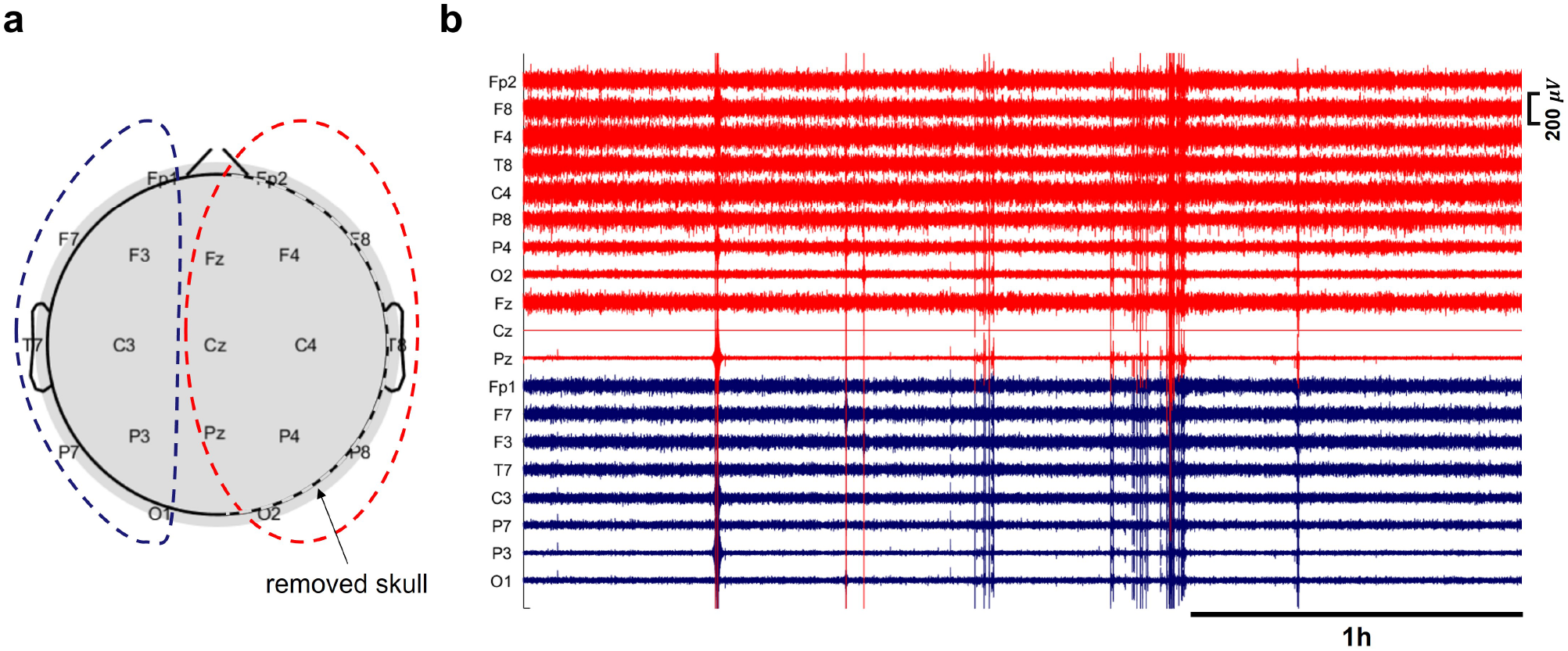
EEG baseline power in ipsilateral (with missing skull) and contralateral (with intact skull) hemispheres, in a patient (patient 4, see Table I for details) with right decompressive hemicraniectomy (DHC): a) 11 ipsilateral EEG electrodes are marked with red dashed line, and 8 contralateral electrodes are marked with blue dashed line, b) 3 hours of the EEG recording across ipsilateral (red traces) and contralateral (blue traces) electrodes. Most of the ipsilateral electrodes on top of the DHC region (with missing skull) have higher EEG baseline power (e.g., Fp2, F8, F4, T8, C4, and P8), in comparison with the contralateral electrodes on the regions with intact skull. The EEG signals are band-pass filtered in [0.5,30] *Hz*, and preprocessed (before amplitude outlier removal, see Section II-C for more information). The signal at Cz had poor quality and was removed through the preprocessing steps.

### C. EEG preprocessing pipeline

We use a multi-step preprocessing pipeline, described below, to prune the continuous EEG recordings and reject ICU-related artifacts and segments of EEG signals with poor quality electrode-scalp contacts:

#### Band-pass filtering and downsampling

We preprocess the EEG data using the EEGLAB toolbox [46] in MATLAB. To be able to evaluate the performance of our SD detection algorithm in different frequency bands, we bandpass filter the EEG signals in different frequency ranges, namely, [0.01, 0.1] *Hz* (near-DC), [0.5, 4] *Hz* (Delta), [4, 8] *Hz* (Theta), [8, 12] *Hz* (Alpha), and [12, 30] *Hz* (Beta) using a Hamming-windowed sinc finite impulse response (FIR) filter. An upper cutoff frequency of 30*Hz* is used to remove high-frequency noise components. The filtered EEG signals are then downsampled to 64*Hz*. We also bandpass filter the ECG and PLETH signals in the frequency range of [0.5, 30] *Hz* and downsampled them to 64*Hz*.

#### “Masking out” poor-quality segments of EEG signal

There are segments of the EEG recordings with poor quality electrode-scalp contacts which could be due to movements of patient on the bed, and conductive gel/saline drying out at each electrode, etc. *CNS Advanced ICU EEG Amplifier* monitors the quality of each electrode-scalp contact through continuous impedance recording at a sampling frequency of 1*Hz*. To enable our algorithm to detect SDs even when a few of the electrodes do not have good contact, we use these impedance recordings to implement “masks” for automated removal of the parts of EEG signals with poor quality EEG. We upsample the continuous recording of impedance at each EEG electrode to match the sampling frequency of the EEG signals. For each electrode, the median of impedance over the whole recording is calculated 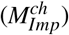 and parts of the EEG signals with abnormally high 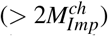 impedance are used for masking. Instead of cutting these parts out of the EEG signals, we assign dummy zero values to these parts (i.e., we “mask out” these parts) to maintain the continuity of the recordings. As it is explained in Section II-D, these *masked-out* sections are excluded from the power envelope extraction and the downstream analyses to prevent false alarms resulting from these zero values. This helps us to keep the parts of the dataset where a few of the electrodes have good recording quality. In Section II-D, we discuss in detail how we detect SDs when the recordings of some EEG electrodes are not available/usable. For the time intervals during which the signals of all EEG electrodes are masked out, dummy zero values are assigned to the PLETH and ECG signals as well for performing independent component analysis, discussed next.

#### Artifact classification and removal using Independent Component Analysis (ICA)

We group together the ECG, PLETH, and EEG signals and perform an ICA to extract and remove sources of artifact (such as eye blinks, eye movements, heartbeats, and muscle artifacts) in EEG signals. We use the EEGLAB [46] toolbox to calculate the independent components and use an automated EEG independent component classifier plugin (ICLabel) in EEGLAB to guide our decision on which components belong to sources of artifacts and subsequently remove them from the EEG recordings.

#### Outlier removal

Extracted ICA components may not perfectly separate some artifacts with abnormally high amplitudes (i.e., outliers) from the EEG signals. Therefore, as the last step in the preprocessing pipeline, we detect and remove the amplitude outliers using Tukey’s fences method [47, 48]. Tukey defines the outliers as data points that fall outside an interquartile range of [*Q*_1_−*k*(*Q*_3_−*Q*_1_), *Q*_3_ + *k*(*Q*_3_−*Q*_1_)], where *Q*_1_ and *Q*_3_ are the first and third quartiles respectively, and *k* is an outlier threshold. We use *k* = 3, which detects “far out” data points according to Tukey’s outlier definition. For each electrode, we detect and mask out (following the same procedure as impedance-based masking discussed earlier in this section) parts of the EEG signals with outlier amplitude. Fig. 3 illustrates the preprocessing steps applied on the ipsilateral EEG signals of a patient with right DHC, in a 4-hour time window. Fig. 3a shows the full-band EEG signals, with poor quality (high impedance) in the first ∼77min of the recording, which is “masked out” in the band-pass filtered signals (in Delta band, see Fig. 3b). The artifacts and outliers are detected and removed as it is shown in Fig. 3c. The preprocessed continuous EEG signals were then used for SD detection using WAVEFRONT.

**Fig. 3.**
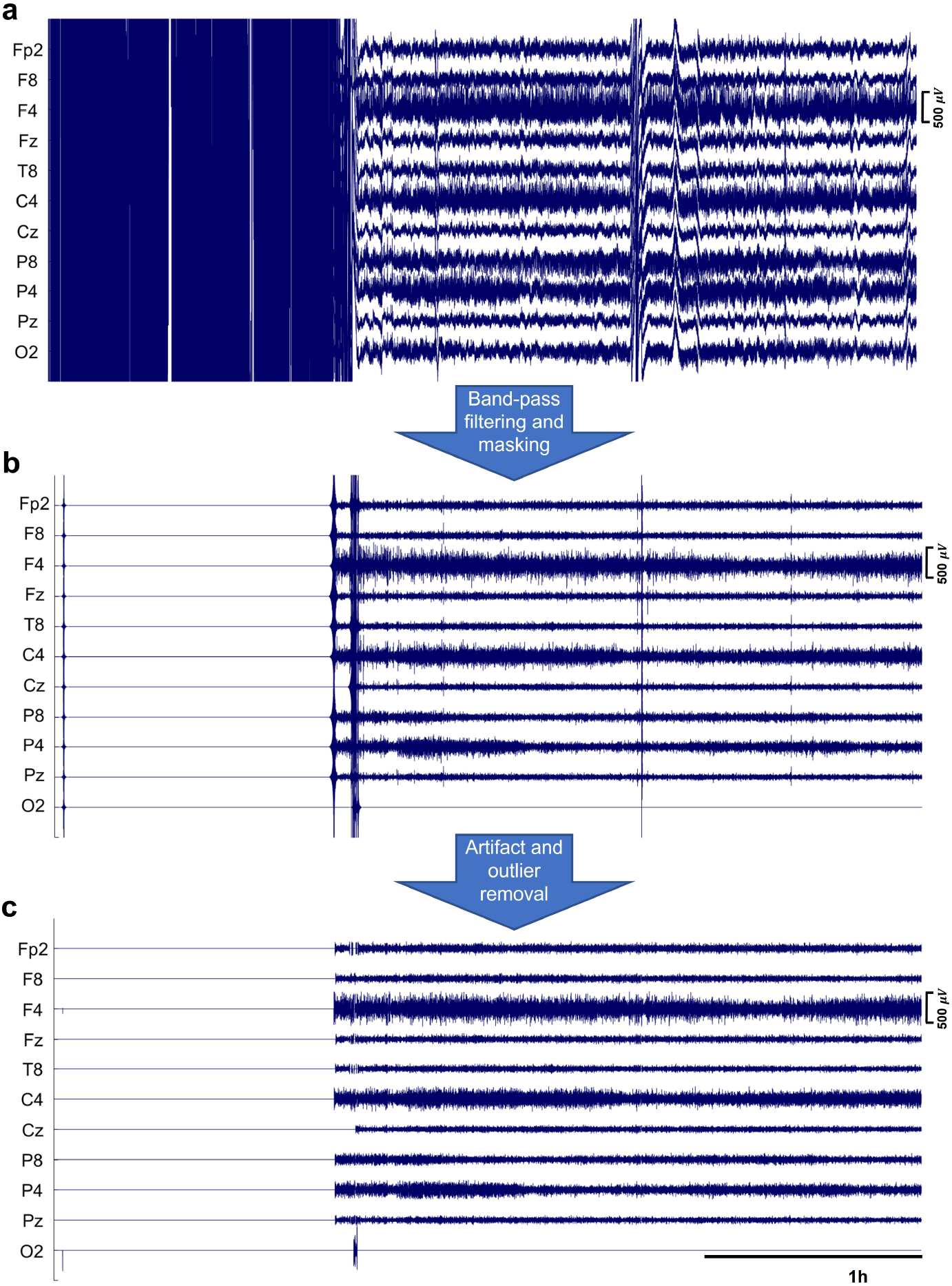
Application of the preprocessing steps on the ipsilateral EEG recordings of a patient with right DHC, in a 4-hour time window: a) the full-band EEG signals, with poor quality (high impedance) in the first ∼77min of the recording, which is “masked out” in the band-pass filtered signals (Delta band, [0.5,4] *Hz*) shown in (b). The artifacts and outliers are then detected and removed in (c).

### D. SD detection and tracking using WAVEFRONT

We use our previously proposed WAVEFRONT SD detection algorithm [28], with appropriate modifications and improvements, to detect and track SD waves in EEG recording of 12 TBI patients in the dataset (see Table I for more details on these patients). WAVEFRONT is an *explainable* automated SD detection framework with intuitive steps and *interpretable* detection outputs and results. It addresses the challenges of noninvasive detection of SD waves in EEG (see Section I for details), by using a computer vision technique, called optical flow, to estimate the speed and direction of propagation of depression wavefronts in the scalp. These depression wavefronts are obtained based on the extracted power envelopes of the EEG signals on the scalp, which are then projected onto a 2D plane for optical flow estimations. Instead of dealing with the challenging task of recovering and estimating the whole underlying propagating SD wave in the brain using the noisy and low-pass filtered scalp EEG signals, WAVEFRONT detects and classifies disjoint SD wavefronts, and stitches together these detections over time and space to detect and track SD waves in the brain. WAVEFRONT suffers from certain shortcomings and cannot be directly applied on real scalp EEG signals for SD detection: (i) it uses fixed and manually adjusted set of parameters (e.g., depression level/depth threshold, temporal score threshold, and spatio-temporal neighborhood radius), (ii) it is highly sensitive to the amplitude outliers, (iii) makes an implicit assumption that the baseline EEG power (i.e., power level of normal background brain activity in the EEG signals) is homogeneous across time and electrodes (see Fig. 9 and 10 in [28] for more details), which limits the ability of WAVEFRONT in detection of depressions in electrodes with low baseline power (e.g., in energy-compromised regions such as ischemic tissue or near lesions), (iv) the depression extraction step is not designed to detect near-DC shifts during propagating SDs in the EEG recordings, (v) does not address the challenges of using a very low density EEG grid, including the high rate of false alarms due to the non-propagating depressions on the scalp, and (vi) estimates the optical flows in pixels on the 2D images, rather than in terms of the physical distances on the scalp.

In this work, we address these limitations in WAVEFRONT by making appropriate modifications and improvements, including designing a training and validation framework for the algorithm to learn an optimal set of parameters, using a rigorous and automated preprocessing pipeline for outlier rejection and pruning the EEG signals (see Section II-C for details), using a power envelope extraction method which is less sensitive to the amplitude outliers, extending the depression extraction method to be able to detect the DC shifts in the near-DC components ([0.01-0.1] *Hz*), as well as the power depressions in the higher frequency bands (≥ 0.5 *Hz*), and defining an “effective propagation measure” along with a learnable threshold on this measure to reject the false alarms of the non-propagating depressions on the scalp. In this section, we include the steps of the WAVEFRONT algorithm, shown in Fig. 4, and provide details on the modifications and improvements we make in each step:

**Fig. 4.**
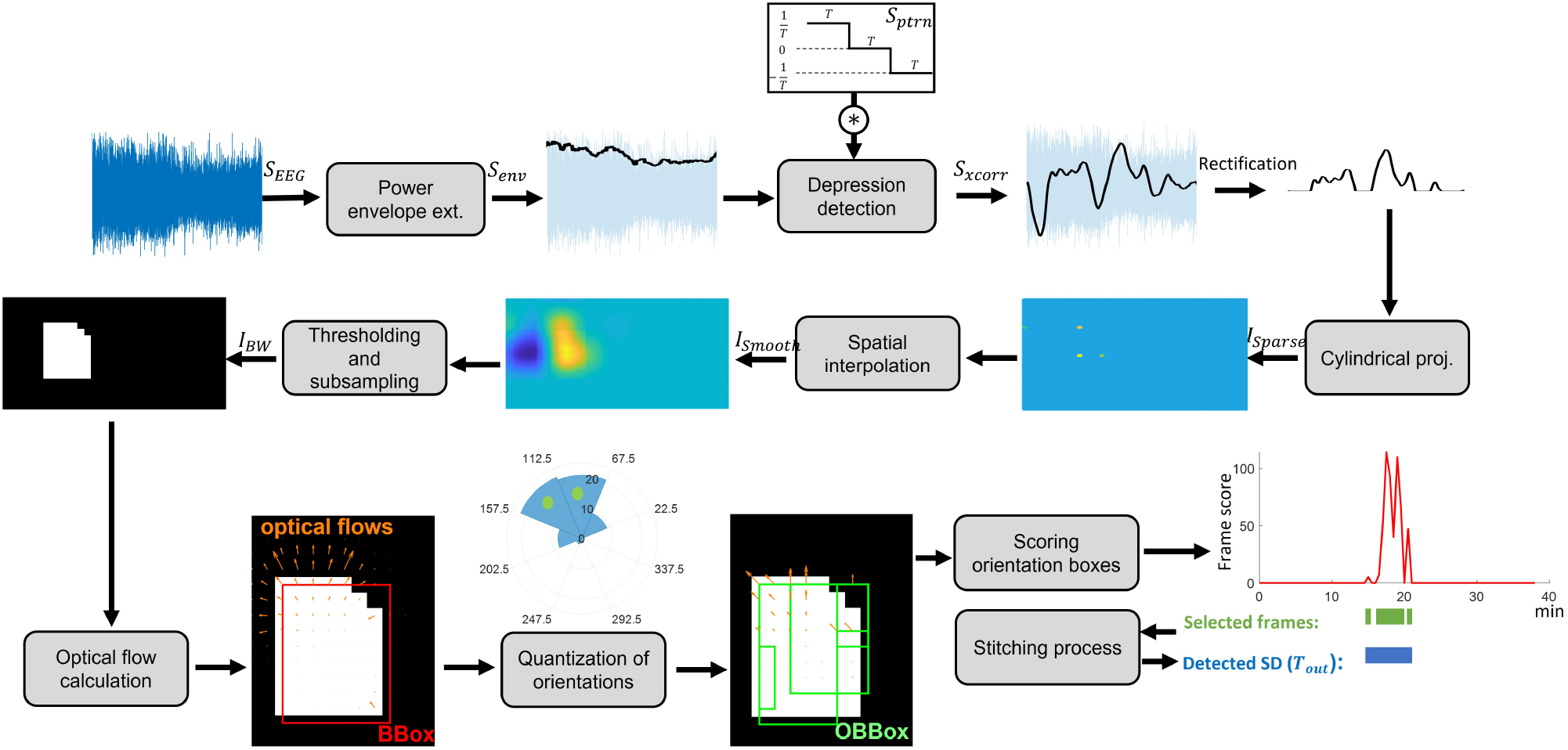
Main steps of the WAVEFRONT algorithm: the power envelope of the preprocessed EEG signals (*S*_*EEG*_) are extracted, and crosscorrelated with a first-derivative kernel (*S*_*Ptrn*_)) to extract power depressions as large positive peaks in *S*_*Xcorr*_, which are rectified and projected on a 2D plane through a cylindrical projection. The resulting image (*I*_*Sparse*_) is then spatially interpolated, thresholded, and subsampled to obtain binary images (*I*_*BW*_), where the depression wavefronts are captured as white contiguous pixels. The movement of wavefronts in these binary images are estimated using optical flows, dominant directions of propagation (marked bins in the orientation histogram) are found through quantization of orientations, and scored based on the consistency and speed of propagation of wavefronts. Candidate frames are selected based on the calculated scores and stitched together for the final detection output (*T*_*out*_).

#### Epoching and envelope extraction

We extracted epochs from the preprocessed EEG signals using overlapping time windows of 240min with a step size of 180min. For each epoch, the EEG signal at each electrode (*S*_*EEG*_) was normalized by its estimated standard deviation. As mentioned in Section II-B, we only use ipsilateral scalp EEG electrodes for each patient because of the missing spatial SD ground truth in the contralateral hemisphere, and heterogeneity of the baseline EEG power between the hemispheres with DHC and the hemisphere with an intact skull. However, even in the hemisphere with DHC, the scalp electrodes which are far from the site of surgery have overall lower baseline power, in comparison to the electrodes which are placed right on top of the regions with a missing skull (see Fig. 2b as an example, where P4, O2, and Pz have smaller baseline power in comparison with the rest of ipsilateral electrodes shown in red). This epoching and power normalization step addresses the heterogeneity of the baseline EEG power across electrodes and over time and helps to detect and extract the depressions in electrodes with low baseline power, which are otherwise missed in the interpolation and thresholding step (this step is explained later in this section). In addition to the baseline power normalization, this epoching helps in the parallelization of the downstream data analysis and the training process explained in Section III-B.

Following the normalization step, the amplitude values are squared, and upper root-mean-square (RMS) envelopes of the power signals are extracted using a sliding time window of 5min. We use the implementation of the RMS envelope extraction method in [49]. There might be some small and isolated valid portions of the EEG signals at each electrode with the normal recording quality, which are interleaved by dummy zero values following the *masking out* step in Section II-C. These portions are not large enough to capture the slow depressions of SDs across EEG electrodes. To prevent false alarms resulting from these isolated intervals, at each epoch and before power envelope (*S*_*env*_) extraction, we *mask out* small intervals of the EEG signals which are less than 20min long, and isolated, i.e., more than 1min apart from the nearest valid intervals. The *masked-out* sections of the EEG signals are excluded from the envelope extraction.

#### Power depression extraction

In this step, we detect and extract the power depressions at each electrode based on the power envelopes (*S*_*env*_). In order to detect the falling edges of the power envelopes, which are followed by a prolonged power reduction, we cross-correlate *S*_*env*_ with a piecewise-constant function as a first-order derivative kernel (see *S*_*Ptrn*_ kernel in Fig. 3). This kernel extracts EEG power depressions as large positive values in the cross-correlated signal (*S*_*Xcorr*_). The 5min width used for envelope extraction and depression edge detection is small in comparison to the large temporal width of depressions in severe TBIs. The same first-order derivative kernel is used for the detection of DC shifts in the near-DC components ([0.01, 0.1] *Hz*). This kernel is directly applied on *S*_*EEG*_, after the epoching and power normalization step, to detect the falling edges of SPCs in the near-DC components. Due to the low density of EEG electrodes on the scalp (only 19 electrodes), Laplacian spatial filtering [50, 51] is not effective to extract narrow SDs [28, 50], and hence we did not use it in this study.

#### Cylindrical projection

We closely follow the steps in [28] to project the extracted *S*_*Xcorr*_ signals at ipsilateral EEG electrode locations on a 2D plane. Before this projection, *S*_*Xcorr*_ signals are *rectified*, i.e., negative values (corresponding to the rising edges in the power envelopes) are zeroed, and positive values, which correspond to the falling edges of the depressions, are kept unchanged. Fig. 4 shows an example of the resulting image (*I*_*Sparse*_) using this projection for patient 3, around an annotated SD event. The corresponding scalp electrode locations in these 2D plane are shown in Fig. 5 for patients with left and right DHC. At each time point, the median value of the *S*_*Xcorr*_ signals at ipsilateral electrode locations is assigned to the rest of the pixels in *I*_*Sparse*_.

**Fig. 5.**
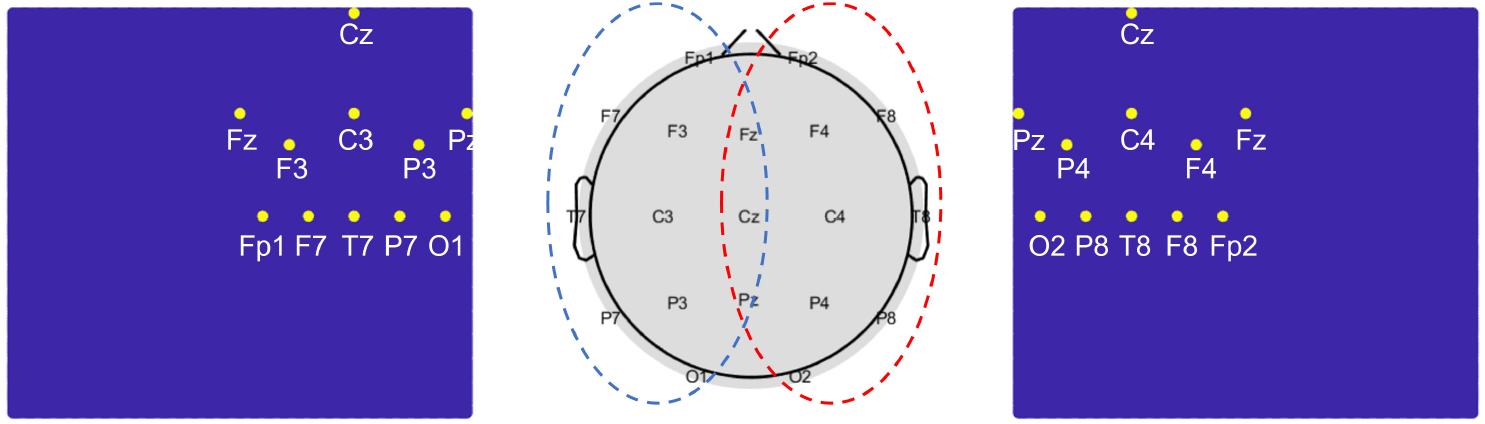
Ipsilateral scalp electrode locations in the 2D images, resulting from the cylindrical projection step in WAVEFRONT (e.g., *I*_*Sparse*_, *I*_*Smooth*_, and *I*_*BW*_), for patients with left (left panel) and right (right panel) DHC.

#### Interpolation and thresholding

We spatially interpolate *I*_*Sparse*_ using a 2D Gaussian kernel with *σ* = 2.62 cm. This large standard deviation was chosen because of the low density of EEG electrodes, where the average inter-electrode distance is ∼5.4cm. For interpolation at the boundaries, each *I*_*Sparse*_ image is padded with the median of the corresponding ipsilateral values of the *S*_*Xcorr*_ signals. The resulting smooth image (*I*_*Smooth*_) is shown in Fig. 4. Following this step, we introduce a thresholding mechanism to extract binary images from the smooth 2D images (*I*_*Smooth*_): (i) we assign zeros to the pixels at each image whose values are 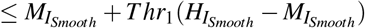 where 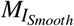 and 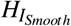*>*are the median and the maximum of the pixel values at each *I*_*Smooth*_ image respectively, and *Thr*_1_ is the depression-level threshold at each time point which are found through the training process as a learnable parameter (see Section III for details on the training and validation steps). This step zeros the pixels with values close to or less than the median value at each image, and set the pixels with large positive values in *I*_*Smooth*_ as candidates for *SD wavefronts*, (ii) In addition, we assign zeros to the whole images where most of the EEG electrodes are masked out (images with less than 5 participating scalp electrodes out of the total 11 ipsilateral electrodes). This rejects the binary images with poor EEG signals, (iii) we set the remaining pixels (assign 1’s), and finally reject (assign zeros) the images where more than half of the pixels are set. This is done with the reasonable assumption that SD depressions cannot spatially expand over more than half of the cortical surface. Following this three-stage thresholding mechanism, binary images (*I*_*BW*_ in Fig. 4) were extracted, where non-zero pixels form connected components which are either parts of the SD wavefronts or non-SD activities in the brain. Through the following steps, we classify these connected components and detect and track SD wavefronts.

#### Subsampling and optical flow calculation

We temporally subsample the series of binary images (*I*_*BW*_) every 30sec. Since the extracted power envelopes and depression signals (*S*_*Xcorr*_) have a very slow temporal pattern in the order of minutes, this temporal subsampling will significantly reduce the computational complexity of the WAVEFRONT for noninvasive detection of SDs in these TBI patients, without adversely affecting the SD detection performance. In addition, due to the very low density of the CNS EEG grid in this study (only 19 electrodes), we spatially subsample *I*_*BW*_ images so that the inter-electrode distances in these 2D images are less than three pixels. This helps to better capture the propagation of SD wavefronts across EEG electrodes in these binary images and further reduces the computational power required for the downstream analysis in WAVEFRONT. We use the bicubic interpolation method [52], and its Matlab implementation [53], for spatial subsampling of *I*_*BW*_. After these subsampling steps, we closely follow the steps in [28] for the calculation of optical flows to capture the movements of connected components and SD wavefronts.

Advancing on [28], we make minor modifications and improvements to the way we estimate optical flows. Optical flow is a computer vision technique to track moving objects across frames of a video [43, 44]. It uses the spatiotemporal brightness variations of the pixels to estimate the velocity (magnitude and direction) of the moving objects. The optical flows of the depressions are calculated based on the 2D images, which are the cylindrical projections of scalp electrode locations onto a 2D plane. In these 2D images, the horizontal dimension captures the azimuthal angle (*ϕ*), and the vertical dimension captures the polar angle (*θ*, with 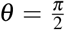 to be at the north pole) of the electrode locations in the spherical coordinate. Therefore, to estimate the optical flows of the connected components (the contiguous non-zero pixels in *I*_*BW*_) in spherical coordinates, based on the calculated optical flows in the 2D images, we define the following mappings for the vertical and horizontal optical flow magnitudes:

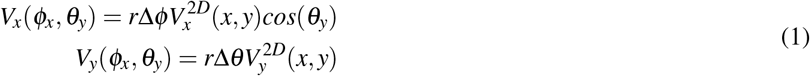

where *r* is an average human head radius (we used *r* = 75*mm* in this study), 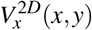 and 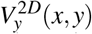 are the horizontal and vertical magnitudes of the estimated optical flow at (*x, y*) on the 2D plane, and *V*_*x*_(*ϕ*_*x*_, *θ*_*y*_) and *V*_*y*_(*ϕ*_*x*_, *θ*_*y*_) are their corresponding magnitudes on the spherical model of the scalp. (*ϕ*_*x*_, *θ*_*y*_) is the corresponding spherical coordinate of the (*x, y*) location in the 2D plane, and Δ*ϕ* and Δ*θ* are the azimuthal and polar resolution of each pixel in the 2D images. Fig. 6 shows an example of this mapping for an optical flow. We use the mapped **V** = (*V*_*x*_,*V*_*y*_) spherical optical flows instead of the original 2D optical flows 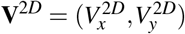 throughout the next steps of WAVEFRONT. It is worth mentioning that **V** = (*V*_*x*_,*V*_*y*_) is an estimation of the extent of movements for the connected components in *mm* on the spherical scalp surface rather than in pixels on the 2D images. This makes it easier to impose constraints on the speed of propagation of connected components for the detection of SD wavefronts (see the “Scoring OBBoxes based on the consistency of propagation” subsection for more details).

**Fig. 6.**
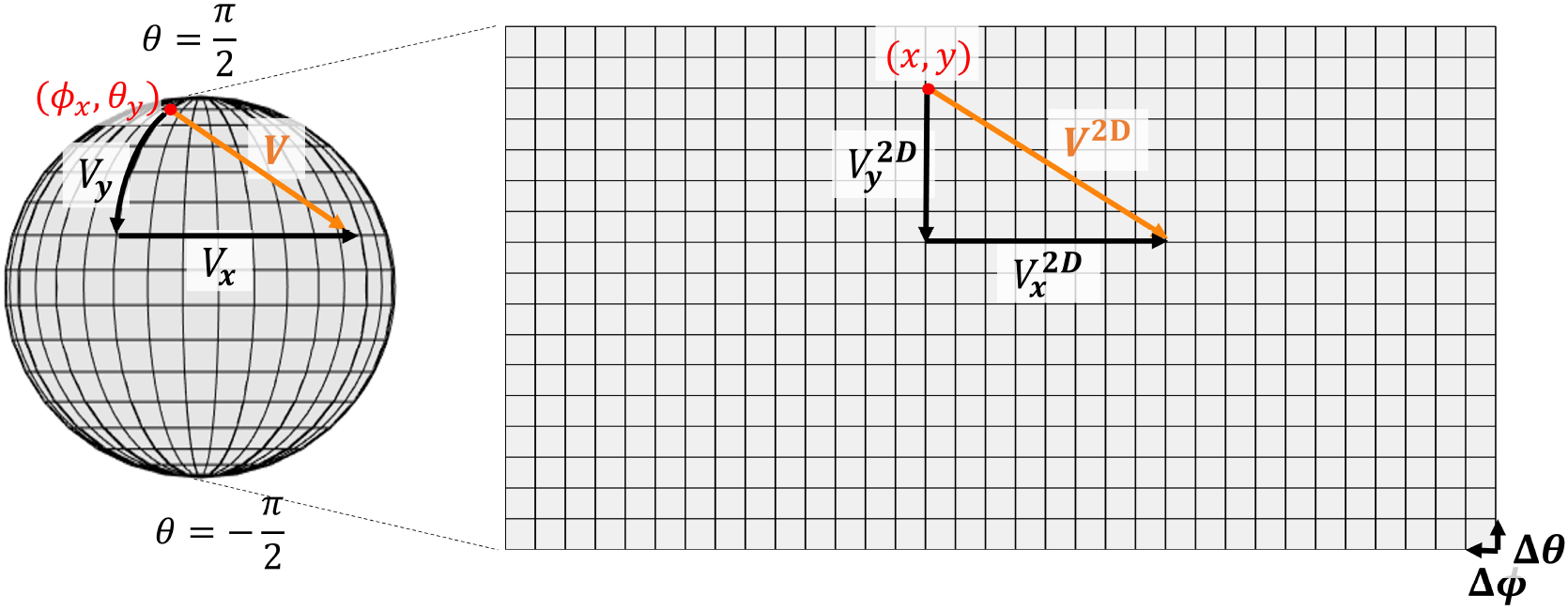
Mapping of an optical flow (**V**^2*D*^(*x, y*)) in the 2D plane on the scalp spherical surface (**V**(*ϕ*_*x*_, *θ*_*y*_)).

#### Quantization of orientations

We closely follow the steps in our previous work [28] to assign bounding boxes “BBox” to the connected components in *I*_*BW*_, calculate an orientation histogram for each BBox and quantize the orientations of optical flows, and finally extract prominent direction(s) of propagation in each BBox. In this study, we introduce an additional constraint on the *effective propagation* of each BBox before quantization. This additional step is designed to reject the “pop-up/fade” types of transition of the connected components in the binary images around each electrode location in *I*_*BW*_. Fig. 7a shows an example of “pop-up/fade” transition in the binary images. The low density of the EEG grid in this study makes it impossible to capture small movements of SD wavefronts unless they propagate sufficiently across the 2D planes. Therefore, to reduce the false alarms because of these “pop-up/fade” transitions, and only considering the propagating depressions across scalp electrodes, we try to detect non-propagating BBoxes and remove them. We define and calculate the “effective propagation measure” or EPM for each BBox, based on the estimated optical flows, as follows:

**Fig. 7.**
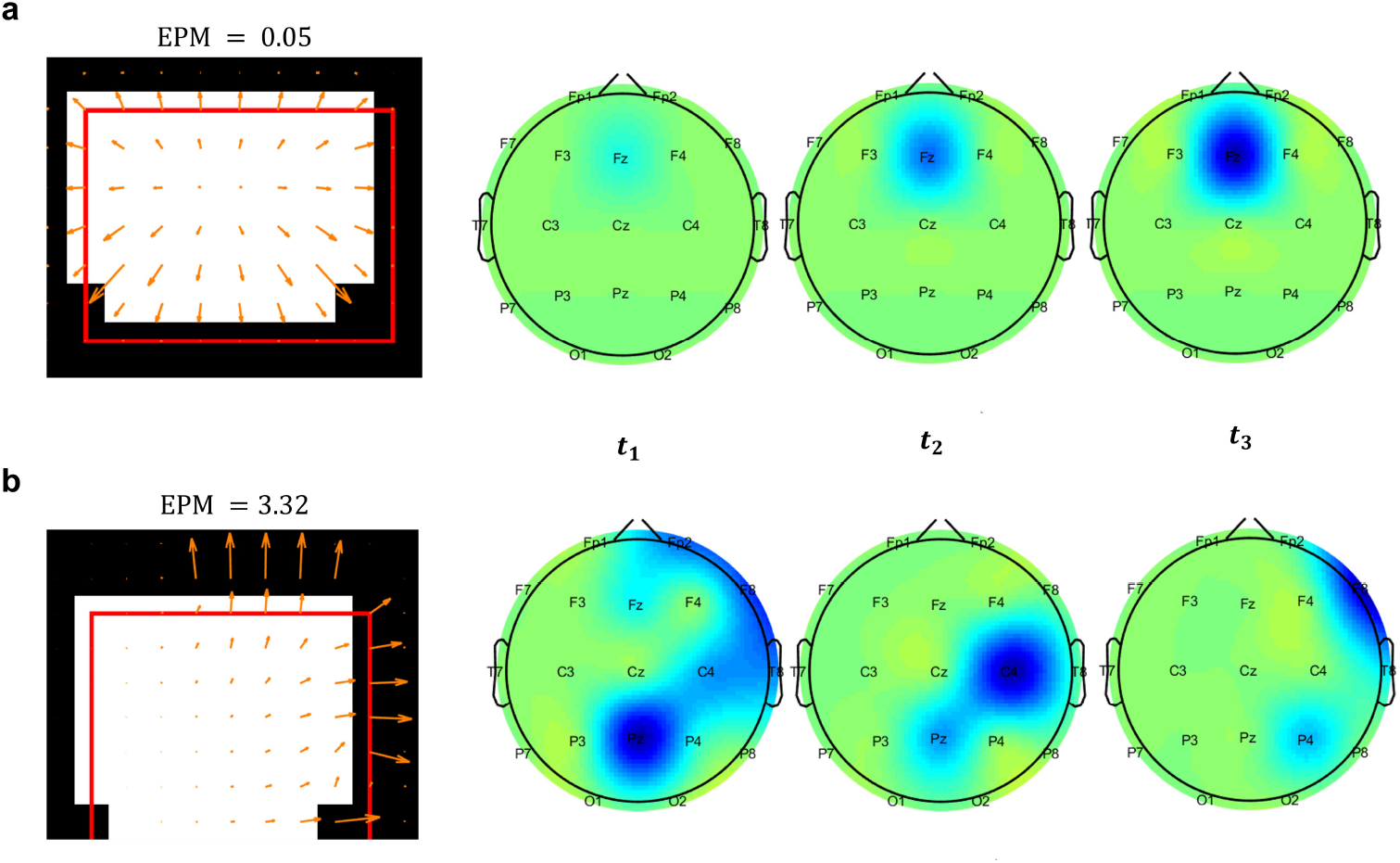
Effective propagation measures (EPMs) in a) a BBox (red box) with a “pop-up” transition (EPM= 0.05), and b) a BBox with significant effective propagation (EPM= 3.32), along with their corresponding depression transitions on scalp (dark blue spots in the scalp topography) at three time points.

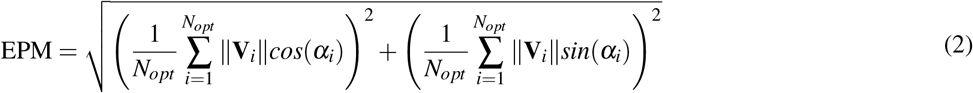

where, ∥**V**_*i*_∥ and *α*_*i*_ are the magnitude and orientation of the *i*^*th*^ optical flow in the BBox, and *N*_*opt*_ is the total number of optical flows in that BBox. The first term under the square root in (2) captures the average horizontal magnitude, and the second term captures the average vertical magnitude of the flows in a given BBox. *EPM* can take values between 0 and 1, where *EPM* = 0 indicates that all optical flows in the BBox are directed either inward (a fading connected component), outward (a popping-up connected component), or have zero magnitudes (no movement in the corresponding connected component). We apply a threshold on *EPM* values of BBoxes and remove the BBoxes and their optical flows if *EPM < Thr*_2_, where *Thr*_2_ is a learnable parameter in the modified WAVEFRONT algorithm (see Section III for details on the training and validation process). The remaining optical flows are quantized using the orientation histograms, following the steps in [28]. Due to the low-resolution EEG grid in this study, we use a coarser orientation histogram with 8 bins of 45^?^ each.

#### Orientations bounding boxes (OBBox)

We extract the orientation bounding boxes (OBBox) using the quantized orientations of optical flows (green boxes in Fig. 4), closely following the steps in [28].

#### Scoring OBBoxes based on the consistency of propagation

We closely follow the steps in our previous work [28] to score the OBBoxes. We have used a spatial radius of 7cm (to cover the large inter-electrode distances in this low-resolution EEG grid), and a temporal range of *Thr*_3_ min (*Thr*_3_ min before and *Thr*_3_ min after the current frame) to find the neighbors of each OBBox. The algorithm learns the temporal range of *Thr*_3_ through the training process. We impose speed constraints on the propagating wavefronts and remove the OBBoxes with very fast (*>*8mm/min) or very slow (*<*0.5mm/min) propagation, count the number of matching OBBoxes for each of the remaining boxes, and consider this count as a “spatiotemporal” score to each OBBox. In addition, we calculate the “temporal score” for each OBBox to measure the consistency in the propagation of wavefronts over time. If the fraction of the number of frames with non-zero temporal scores over the total number of frames in the *Thr*_3_ min neighborhood is less than *Thr*_4_ (i.e., only a small number of frames contribute to the “spatiotemporal” score), we remove the corresponding OBBox (i.e., assign zero to its “spatiotemporal” score). *Thr*_4_ is another learnable parameter in the WAVEFRONT algorithm and takes values between 0 (no contributing frame to the score of the OBBox) and 1 (all frames in the temporal neighborhood of the current frame, defined by *Thr*_3_, contribute to the score of the corresponding OBBox).

#### Stitching process and the final decision on detection

We reject the OBBoxed with small “spatiotemporal” scores (less than 1% of the maximum available score at each frame), and reject the frames with small “frame scores” (scores of less than 5% of the median of the frame scores). As the final step, we stitch together the selected frames using a sliding time window of 2min and closely followed the steps in [28] to obtain the final temporal detection output *T*_*out*_ (1 = detected SD at the corresponding frame, 0 = no SD wavefront was detected at the corresponding frame). Fig. 11 shows an example of SD detection for patient 6, with a single isolated SD event in a ∼3-hour time window. Fig. 10 illustrates an example of SD detection in a patient (patient with highly clustered SDs (8 annotated SD events in ∼3 hours of recordings), where the underdetection of WAVEFRONT is apparent, along with some missed detection intervals. These detection examples are based on Delta band EEG signals, and using an optimal set of parameters in WAVEFRONT (see Section III-B and Fig. 8 for details). Please note that in this study, we do not use the spatial detection output (*I*_*out*_) of the WAVEFRONT algorithm for performance evaluation since we lack the spatial ground for SD wavefronts. Detailed discussions on the limitations of WAVEFRONT and the ground truth annotations are included in Section IV.

**Fig. 8.**
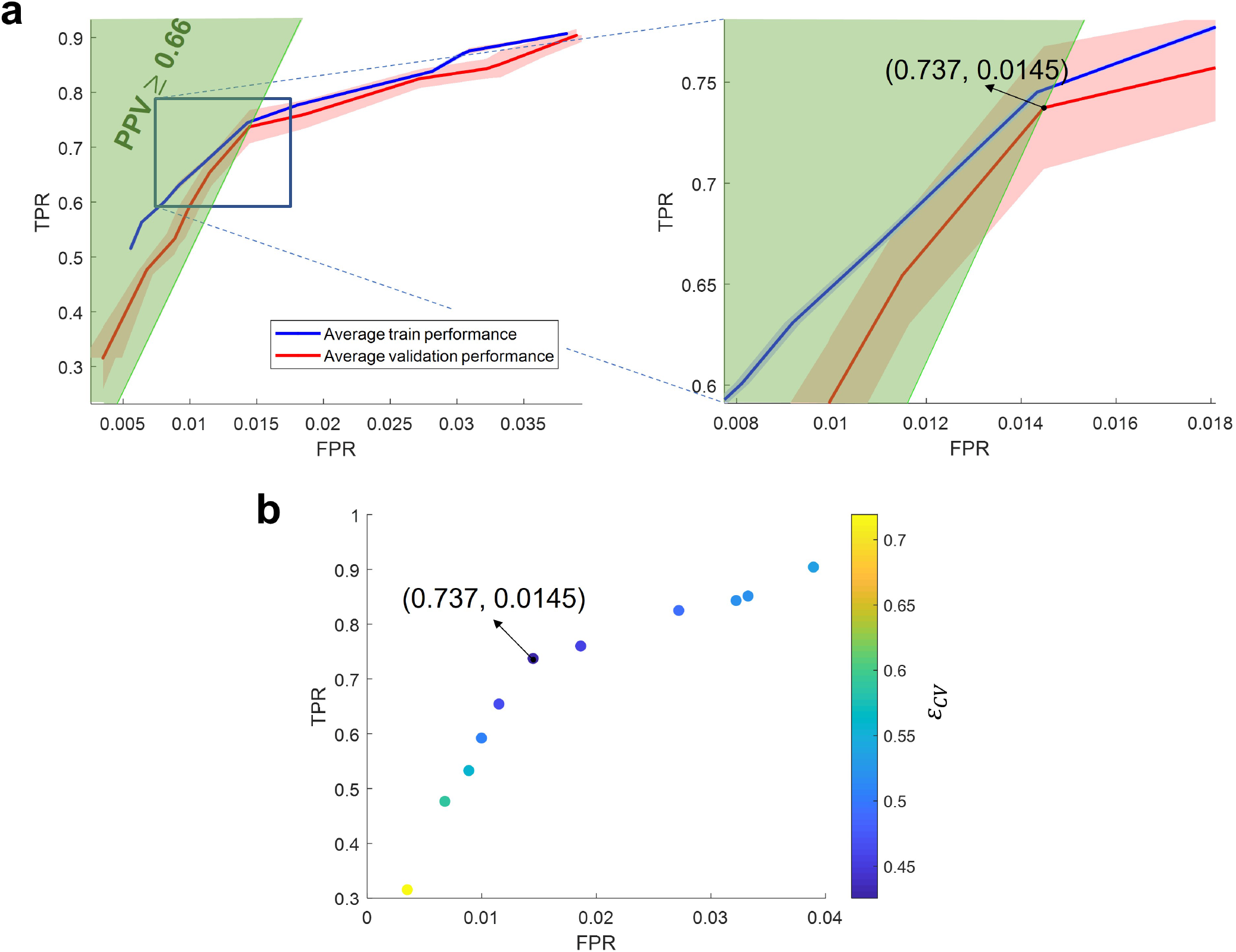
Performance evaluation using a cross-validation analysis: a) Receiver operating characteristic (ROC) curves of the average train (solid blue curve) and validation (solid red curve) performance of WAVEFRONT in the detection of spreading depolarization (SD) events, using noninvasive scalp EEG signals in Delta frequency band. The shaded blue and red regions show the 95% confidence intervals for the average train and validation curves respectively. The vertical axis indicates true positive rates (TPRs), and the horizontal axis shows false positive rates (FPRs). The green highlighted region indicates the positive predictive values of ≥ 0.66 (PPV). The right panel illustrates the zoomed-in version of the ROC curves around the optimal validation operating point (TPR= 0.74±0.03, and FPR= 0.0145±7.57 × 10^−4^, in 95% confidence intervals), b) cross-validation error (*ε*_*CV*_), color-coded across different points in the validation ROC curve, where the point with the minimum error (i.e., optimal operating point) is marked.

## III. RESULTS

In this section, we quantify the performance of our modified WAVEFRONT algorithm, as described in Methods, on 12 TBI patients in the SD-II dataset (see Table I and Section IIA for more details on these patients). We explore the generalizability of our algorithm through a cross-validation analysis and compare the performance of WAVEFRONT across different EEG frequency bands. With an emphasis on the trustworthiness of our method, we provide different visual illustrations of the SD detection results, including temporal and spatial visualization of representative SD events, and carefully define the performance metrics that we have used in this study. Finally, we evaluate the performance of WAVEFRONT in measuring the frequency of SDs (the number of occurrences) in large (30-hour) time windows using regression analysis.

### A. Performance rules and metrics

We assessed the average SD detection performance of WAVEFRONT by examining the performance on overlapping time windows, each with a width of *WL* = 2min and step size of Δ*W* = 30sec. In doing so, we use the following conditions and definitions:

- If a time window includes an annotated SD, it is called an “SD window”. An SD window is said to be *detected* by WAVEFRONT if there exists a non-zero *T*_*out*_ value within a temporal distance of Δ*t* from the annotated SD in the SD window. We choose Δ*t* = 1*hr*, because of the following reasons: (i) for visually observed SDs in DHC patients based on noninvasive EEG signals, the reported time interval between the lowest depression points at two scalp electrodes is 17min (median) with 11-34min interquartile range [22]. Therefore, we examine within Δ*t* = 1*hr* around each SD annotated event, (ii) the average inter-electrode distance of the EEG system used in this study is ∼5.4cm, and the reported range of speed of SD propagation is 1-8mm/min [45]. Consequently, it takes ∼54min for the slowest depression to propagates between each pair of electrodes on scalp, (iii) in addition, the temporal annotation of SD is extracted using the ECoG strip which only covers a local region, while the detection is based on the ipsilateral EEG electrodes which cover the whole DHC hemisphere. Therefore, an SD event may be detected at any time within the duration of propagation, and hence within a temporal distance from its annotation. This is a limitation of the SD ground truth in this study, and the choice of Δ*t* = 1*hr* is only an assumption, which may result in slight over or underestimation of the actual performance of WAVEFRONT in the detection of SD events. A performance metric of true positive rate (TPR) is defined based on the SD windows as follows:

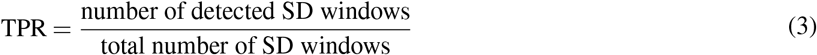

- If a time window includes detection (intervals with non-zero *T*_*out*_), and no annotated SD is found within a temporal distance of Δ*t* from the detected intervals in that window, it is considered a “false alarm” window. In addition, time windows without any detection and any annotated SD inside or within a temporal distance of Δ*t* from either end of the windows are considered “true negative windows”. A performance metric of false positive rate (FPR) is defined based on true negative and false alarm windows as follows:

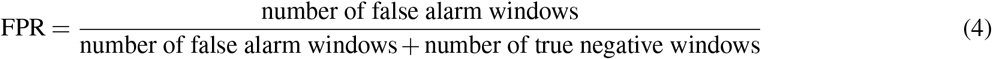

For diagnosis of worsening brain injuries and clinical applications, achieving a low false alarm rate is crucial to minimize the risks and side-effects of unnecessary treatments and interventions, especially invasive interventions such as DHC, for minimizing secondary brain injuries. However, since the dataset is highly imbalanced (the number of non-SD windows is much larger than the number of SD windows), a seemingly low FPR can still have a large number of false alarm windows. Therefore, we use an additional performance metric, defined below.

- We used a third performance metric, called precision or positive predictive value (PPV) [54], defined as:

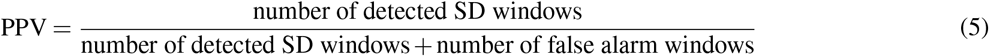

- We also define *Q*_*avg*_, a measure of signal quality for each (sliding, 2 min) time window, as the number of electrodes, averaged over the 2-min interval, that are not masked out in the window over the hemisphere with DHC. Since there are 11 electrodes ipsilateral to the site of ECoG placement (see Fig. 2a, we include electrodes on the midline in the ipsilateral set), we choose a threshold of *Q*_*avg*_ ≥ 6 for defining whether a time-window has good-quality recordings. Thus, time windows with *Q*_*avg*_ *<* 6 are excluded from the SD detection performance calculations. In all, there were 36,709 excluded poor-quality windows (approximately 28% of the windows) across 12 patients. This large number of poor-quality windows is mainly due to the long time intervals during which the patients are disconnected from the EEG amplifier for procedures or imaging. During these intervals, the recordings were not stopped. Other poor-quality intervals may be, in part, due to the inherent limitations of scalp EEG recordings, e.g., low-density of EEG electrodes (only 11 ipsilateral) at ICUs increases the chance of recording intervals with almost no reliable EEG signal. Higher spatial coverage of EEG electrodes on the ipsilateral hemisphere can mitigate this issue. In addition, DHC patients have highly concave scalp surface on the hemisphere with missing skull [35], which makes the scalp electrode placement, and creating a good electrode-scalp contact, even more challenging. This is another important contributing factor for the large number of poor-quality windows in this study. Nevertheless, there is a large number of remaining good-quality SD and non-SD windows (92,583 in total), which are distributed across the 12 patients and used for the calculation of WAVEFRONT’s SD detection performance.

All bounds on the average TPR and FPR performance metrics reported here are 95% confidence intervals, which are estimated using the weighted bootstrapping method [55, 56], with a bootstrap sample size of 100 (randomly selected with replacement). The weights are the number of non-SD windows for FPR confidence intervals, and the number of SD windows for TPR confidence intervals.

### B. Testing generalizability of WAVEFRONT using a cross-validation analysis

#### Leave-2-out cross-validation

To evaluate the generalizability of WAVEFRONT, and detect and prevent overfitting of our algorithm [57], we use cross-validation: we split the dataset into sets of train and validation patient groups, find the optimal sets of parameters for WAVEFRONT on the train sets, assess the SD detection performance on the validation sets, and average the performance on different validation sets. Specifically, we use Leave-2-out cross-validation (L2O CV). We choose two patients out of the total 12 patients, and leave them out for validation, in 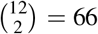 different ways. For each of these 66 choices, we fine-tune WAVEFRONT parameters using the 10 patient-training set, following the steps below:

Using the defined performance metrics in Section III-A, we optimize the parameters 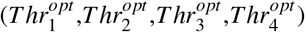 in WAVEFRONT for the best possible train performance, i.e., lowest possible FPR, while maintaining a good detection power (high TPR) and precision (high PPV). This optimized set of parameters is then used to evaluate WAVEFRONT’s performance on the corresponding validation set. We use a brute-force grid search to find the best set of parameters (list of search grids is included in Table II). Following are the training and validation steps for each pair of train-test sets: (i) using the values in the search grids and for each set of (*Thr*_1_,*Thr*_2_,*Thr*_3_,*Thr*_4_), the performance of WAVEFRONT is evaluated on the train set, (ii) a TPR threshold (*Thr*_*TPR*_) is then applied on the train performance, and among the sets of parameters with train TPR≥ *Thr*_*TPR*_, the one with the minimum FPR is chosen as the optimized set, (iii) WAVEFRONT, using the optimized set of parameters, is applied on the validation set to obtain the validation performance. We repeat these steps for all of the train-validation pairs and average the train and validation performance across all of the 66 pairs of train-validation sets. We calculate the average performance for different *Thr*_*TPR*_ values in the range of [0.5, 0.85], and (iv) the optimal operating point (i.e., 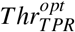), and its corresponding optimal set of parameters, is found using the underfitting-overfitting (also known as bias-variance) <u>tradeoff [58]. We define</u> a cross-validation root-mean-square error (RMSE) using the averaged TPR and PPV values as 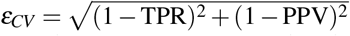. The average validation performance corresponding to the minimum cross-validation error *ε*_*CV*_, with PPV≥0.5, is reported as the generalization performance. PPV= 0.5 is the threshold at which only half of detected intervals are true positives and the other half are false positives.

**Table II.** Search grids for the WAVEFRONT’s parameters. Parameter Search grid

| Parameter | Search grid |
| --- | --- |
| $Thr_1$ | $[0.1, 0.14, 0.18, \dots, 0.9]$ |
| $Thr_2$ | $[0.05, 0.10, 0.15, \dots, 0.8]$ |
| $Thr_3$ | $[2, 3, 4, \dots, 10]$ |
| $Thr_4$ | $[0.6, 0.7, 0.8, 0.9]$ |

The TPR-FPR trade-off is captured in a receiver operating characteristic (ROC) curve. We closely follow the threshold averaging method in [59] to generate the average train and validation curves. Fig. 8a shows the average train (solid blue line) and validation (solid red line) ROC curves of WAVEFRONT performance in detection of SDs using scalp EEG Delta band ([0.5, 4]*Hz*), with TPR values along the vertical axis, and FPR values along the horizontal axis. PPV line of PPV= 0.66 is overlaid on top of the ROC curves, where the PPV*>* 0.66 region is located above the corresponding PPV line. Fig. 8b shows the crossvalidation error (*ε*_*CV*_), color-coded across different points in the validation ROC curve, where the point with the minimum error (i.e., optimal operating point) is marked. Based on the results, using Delta frequency band scalp EEG, WAVEFRONT achieves an average validation performance of TPR= 0.74±0.03 (12,303 of 16,685 total SD windows are detected) in the PPV*>* 0.66 ROC region, with FPR*<* 0.015 (0.0145±7.57 × 10^−4^, 6,339 of 437,849 total non-SD windows are falsely detected). All of the reported results are in 95% confidence intervals. This operating point in the average ROC curve corresponds to an optimal set of parameters as 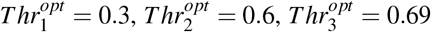, and 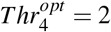. This point has the smallest cross-validation error of 0.4256, while the points with higher TPR show an increasing trend in the cross-validation error (i.e., overfitting [57, 58]), and points with lower TPR have larger error as well (i.e., underfitting [57, 58]). In this study, overfitting is inevitable due to the small number of patients. We expect WAVEFRONT to achieve a better validation performance using a larger data set for the training process. This requires further investigation when we get access to the recordings of more patients with SDs.

Similar TPR-FPR ROC curves for other frequency bands, including the near-DC components, are provided in Fig. 12 (see Section III-C for details).

**Fig. 9.**
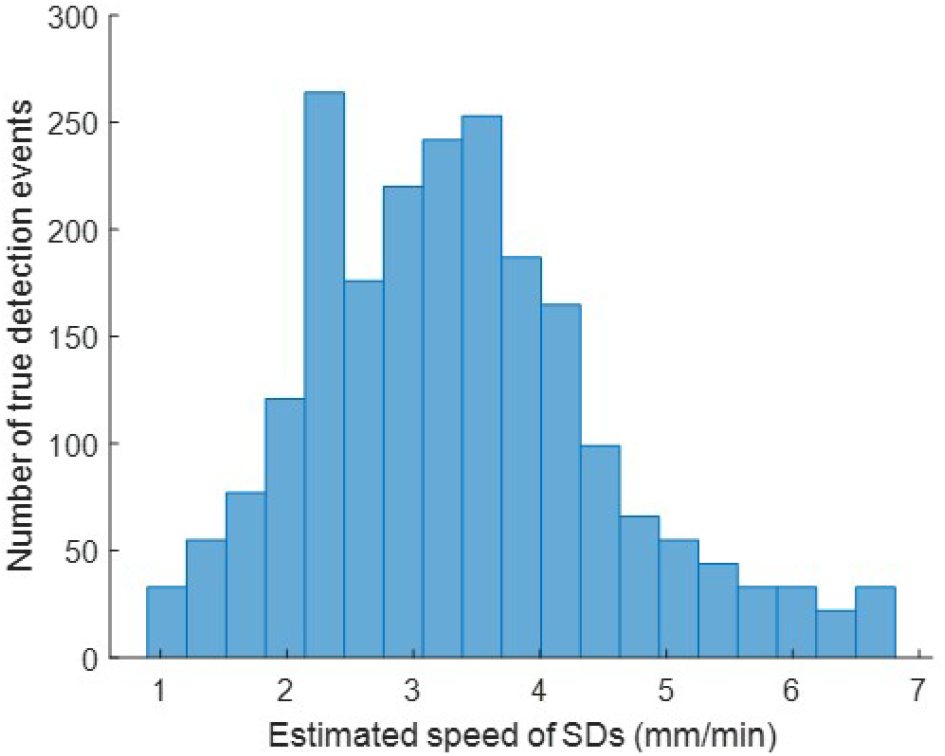
Histogram of estimated speed of propagation of detected SD events in the scalp EEG Delta band. The speed of propagation of SD waves ranges from 0.9 to 6.8 mm/min, with the maximum population around 3.6 mm/min for 12 severe TBI patients in the dataset.

We estimated the speed of propagation of detected SD events at the found optimal performance point in the Delta band. For each true detection event (connected 1’s in *T*_*out*_ which lie within Δ*t* temporal distance of the annotated SD events), we average over the magnitude of optical flows of the corresponding detected OBBoxe, in the scalp spherical coordinates (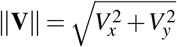 see Section II for more details). Fig. 9 shows the histogram of the estimated speed of propagation for the true detection events in the Delta band. Although no ground truth is available for the speed of propagation of annotated SD events in the dataset, Fig. 9 helps in understanding and comparing the range of SD speeds in the scalp EEG with the available scientific literature. Based on the results, the estimated speed ranges from 0.9 to 6.8 mm/min, which is a slightly smaller range in comparison to the imposed speed constraints in WAVEFRONT ([0.5, 8] mm/min, see Section II), with the largest population around 3.6mm/min. This observation is consistent with the widely reported range of 1-8mm/min in the literature.

Fig. 10 includes a spatio-temporal visualization of a sample SD propagation event in patient 4 with clustered SDs. We order the *S*_*Xcorr*_ time series in Fig. 10b using transverse and longitudinal montages of the EEG electrodes (see Fig. 10a) to make it easier to visually track the propagation of depressions in the extracted *S*_*Xcorr*_ signals (see Section II). In this example, the SD wavefront starts at Fp2 and Fz, and gradually travels towards F4, and ends at T8 and C4 (spatial propagation of this SD wavefront is shown in Fig. 10d). Fig. 10c shows the MRI (left) and CT (right) scans of this patient, where the location of the six ECoG electrodes (right frontotemporal) is shown, along with the locations of lesions and DHC. In this sample visualization in Fig. 10, there are 8 clustered SD events (more than two SDs in a time interval of 3 hours or less [10, 19]), where WAVEFRONT detects 5 SD events in 3 detection intervals (blue strips in Fig. 10b). This illustrates how WAVEFRONT, at least with low-density EEG, can underdetect SD events, especially when they are clustered.

**Fig. 10.**
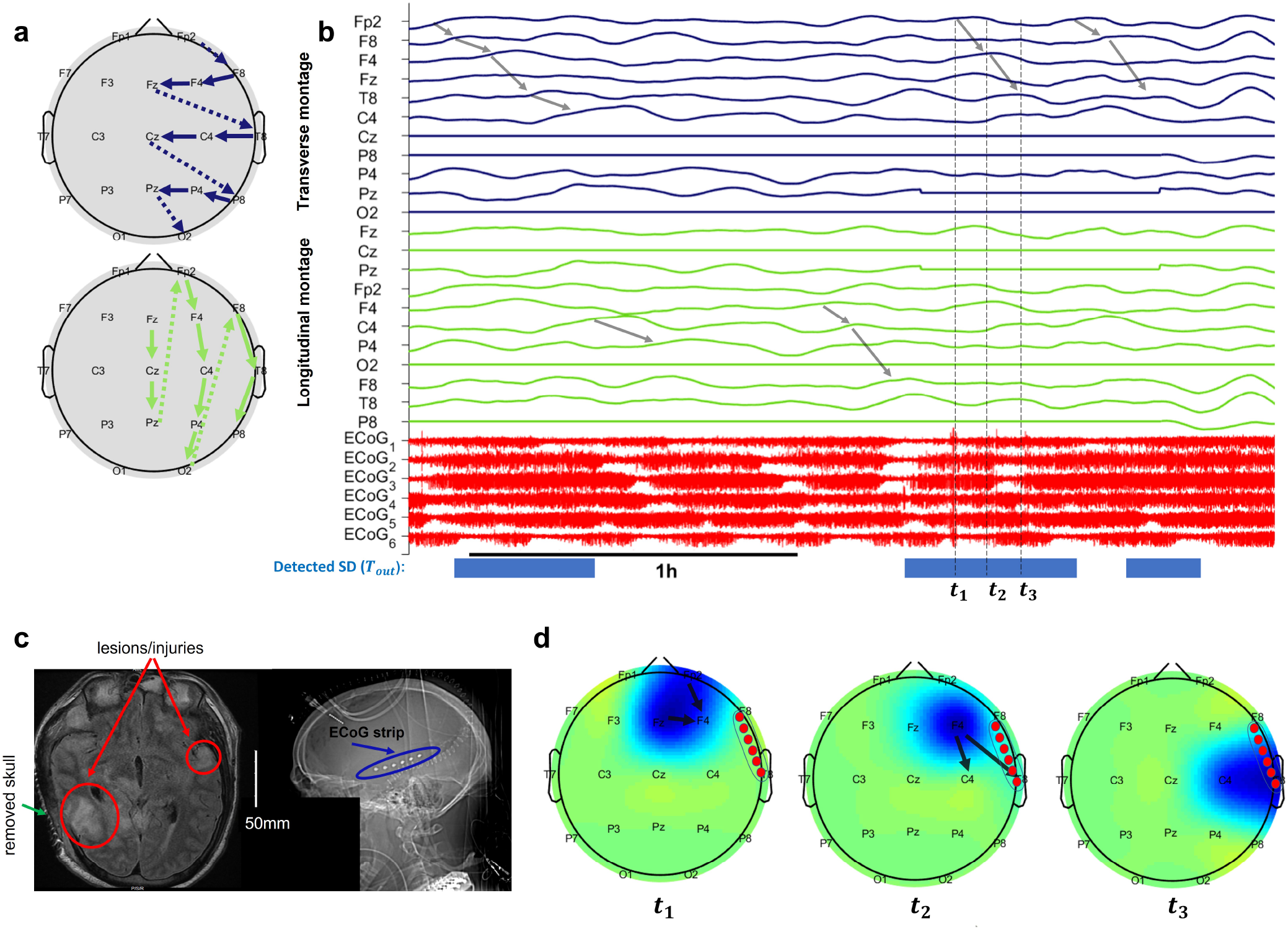
Visualization of a sample SD event in patient 4 with clustered SD events: a) transverse and longitudinal montages of ipsilateral EEG electrodes. These montages order the electrodes in a way so that the signals of anatomically neighboring electrodes are located next to each other in the temporal plots, b) time traces of *S*_*Xcorr*_ and ECoG signals, where three time-points of the selected SD event are marked as *t*_1_, *t*_2_, and *t*_3_ with maximum depressions (peask in *S*_*Xcorr*_) at (Fp2, Fz), F4, and (C4, T8) respectively, c) MRI (left) and CT (right) scans of this patient, where the locations of lesions and injuries are shown, along with the right decompressive hemicraniectomy (DHC) region and the intracranial strip of ECoG electrodes, and d) scalp topography of SD depressions at the three corresponding time points. The intracranial ECoG strip is located around the right frontotemporal lobe. The detected events (*T*_*out*_ = 1) using WAVEFRONT are marked with blue strips in (b), where the underdetection of WAVEFRONT is apparent, along with some missed detection intervals. A total of 5 SD events are detected in EEG, when 8 are marked in the ECoG signals. Some of the propagating depressions in (b) are marked with gray arrows across the *S*_*Xcorr*_ signals, which correspond to the SD events in this time window.

In Fig. 11, a similar visualization is shown for a single and isolated SD event in patient 6 (see the CT scan of this patient in Fig. 1). In this example, the SD depression first shows up at Fp2 on the scalp, and slowly propagates towards F4, and ends at C4 (a longitudinal path), and at the same time, another propagation is observed far away from the ECoG strip (shown as 6 small red circles), following the path of P4-Pz-O2. The observed depression at C4 is spatio-temporally consistent with the intracranially annotated depression at *t*_3_.

**Fig. 11.**
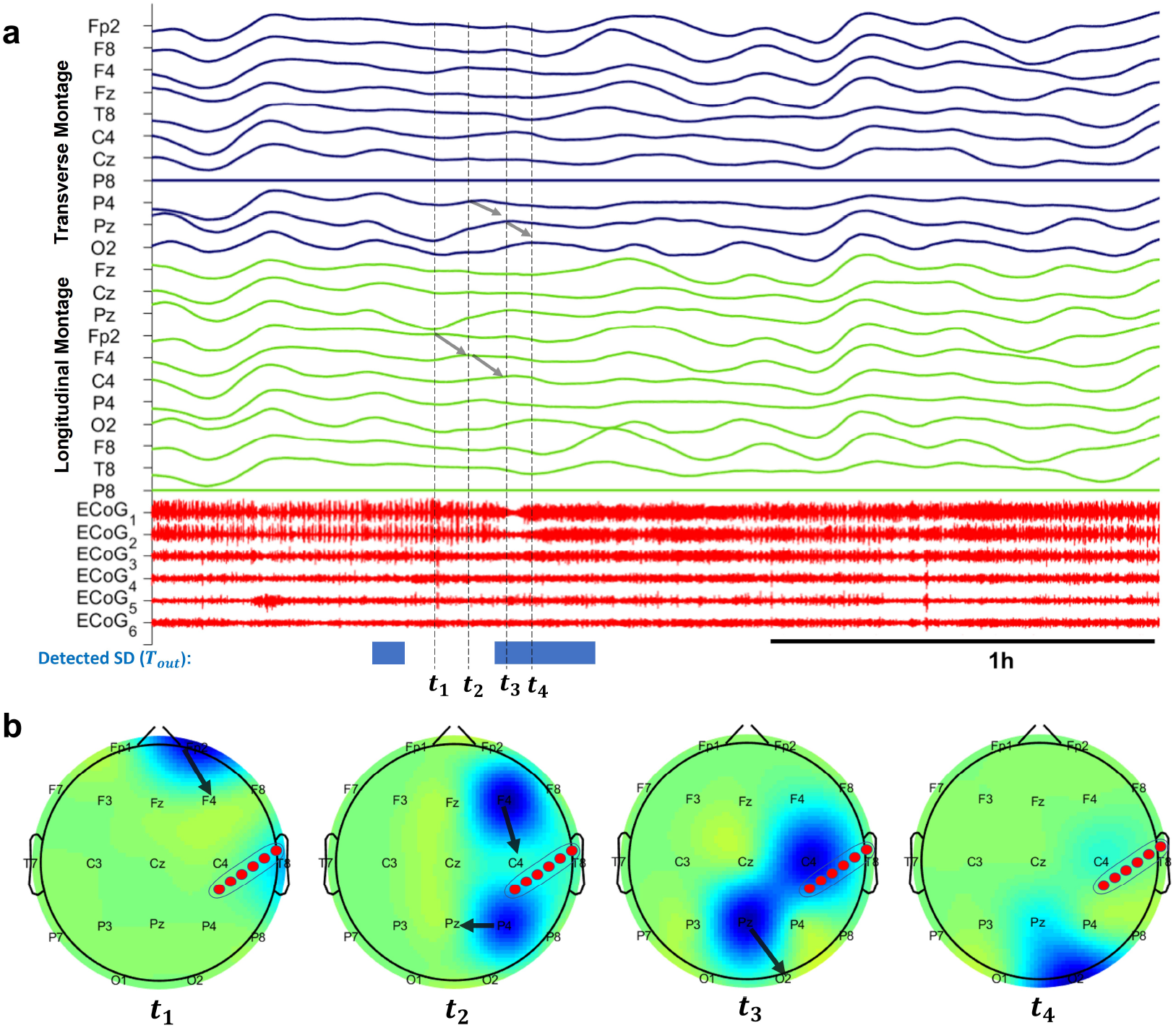
Visualization of a single isolated SD event in patient 6 with right decompressive hemicraniectomy (DHC) (see the CT scan of this patient in Fig. 1): a) time traces of *S*_*Xcorr*_ and ECoG signals, where four time-points of the selected SD event are marked as *t*_1_, *t*_2_, *t*_3_, and *t*_4_ with maximum depressions at Fp2, (F4,P4), (C4,Pz), and O2 respectively, b) scalp topography of SD depressions at the four corresponding time points. The intracranial ECoG strip is located around the right temporoparietal lobe. The detected events (*T*_*out*_ = 1) using WAVEFRONT are marked with blue strips in (a).

### C. SD detection performance using different scalp EEG frequency bands

As discussed in Section II, we band-pass filtered the scalp EEG signals in different frequency ranges of [0.01, 0.1] *Hz* (near-DC), [0.5, 4] *Hz* (Delta), [4, 8] *Hz* (Theta), [8, 12] *Hz* (Alpha), and [12, 30] *Hz* (Beta) to explore the feasibility and performance of noninvasive detection of SD events across frequency bands. An SD propagation may show up as propagating DC shifts across electrodes in the near-DC components, or propagating depressions (power reductions) in higher frequency components (*>* 0.5 *Hz*). In [22], Hartings *et al*. reported that EEG Delta band power, on average, depressed to 47% of its baseline during SD events which are observable in the EEG recordings, where other higher frequency bands experienced less power reduction (i.e., Theta, Alpha, and Beta bands maintained around 60% or more of their baseline power). In addition, since around 81% of the total power of baseline EEG (without SD) is concentrated in the Delta band [22], the contrast between the background EEG power (baseline) and the maximum depressions during SD episodes is much higher for the Delta band in comparison to the higher frequency bands. Therefore, we expect to observe a decreasing trend of WAVEFRONT performance as the function of frequency bands. To test this hypothesis, we train and validate WAVEFRONT based on different frequency bands of near-DC, Theta, Alpha, and Beta, by closely following the steps in Section III-B. The average validation ROC curves for different frequency bands are shown in Fig. 12. Based on the results, in the same ROC region of PPV ≥ 0.50, Delta band achieves the best detection performance (TPR= 0.74±0.03, FPR= 0.015±7.57 × 10^−4^), followed by Theta (TPR= 0.73±0.031, FPR= 0.020±0.0015), near-DC (TPR= 0.70±0.029, FPR= 0.021±0.0011), Alpha (TPR= 0.65±0.039, FPR= 0.020±0.0011), and Beta (TPR= 0.59±0.034, FPR= 0.016±0.0011), all reported in 95% confidence intervals. This SD detection performance trend across different frequency bands is consistent with the expected outcome based on the reported results in [22] using visual inspection of SD events in the scalp EEG recordings. Although the propagating DC shifts are well-known signatures of SD waves in the ECoG recordings of the brain [8], WAVEFRONT has lower SD detection performance using near-DC components of scalp EEG signals (4% less TPR, 0.6% more FPR, and 13% less PPV), in comparison to the best performance among higher frequency bands, i.e., the Delta band. Previous works also reported no propagating DC shifts [9, 23], or very few consistent DC shift propagations in the scalp EEG signals, in comparison to the ECoG recordings of the SD events [22].

**Fig. 12.**
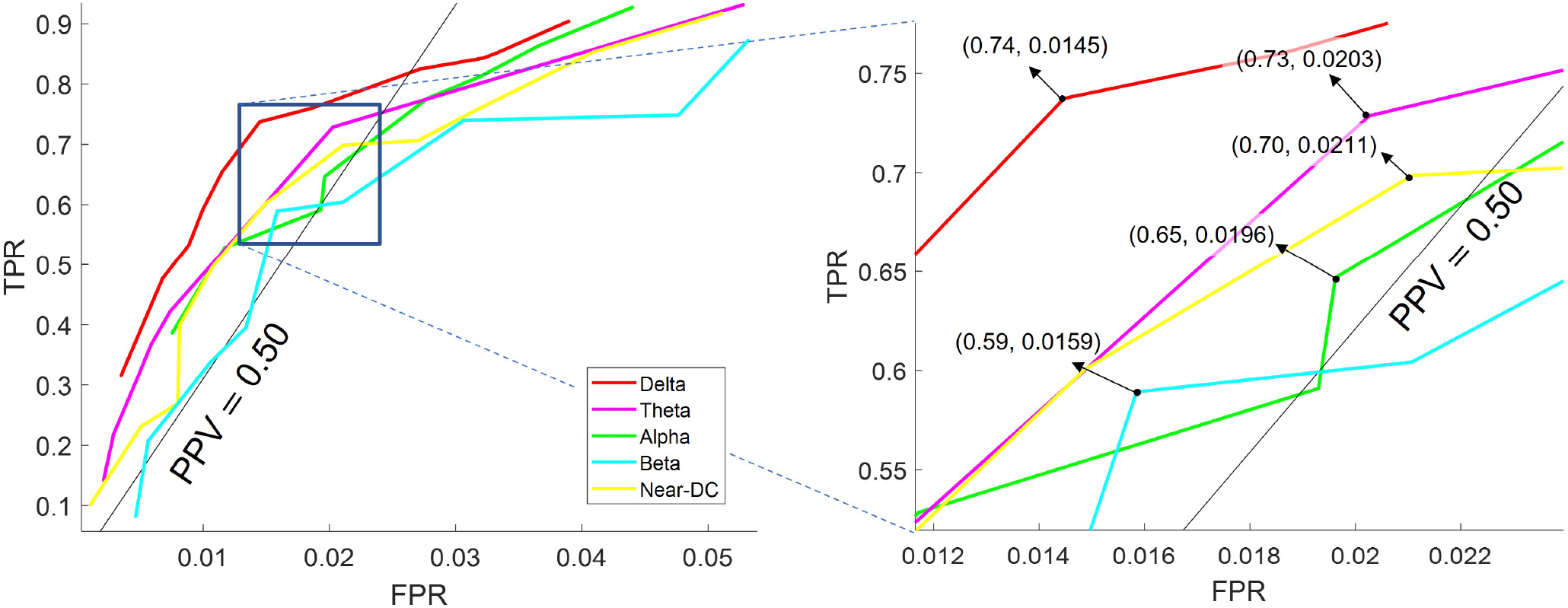
Receiver operating characteristic (ROC) curves of average validation performance of WAVEFRONT algorithm in the detection of spreading depolarization (SD) events, using noninvasive scalp electroencephalography (EEG) signals across different frequency ranges of [0.01, 0.1] *Hz* (near-DC, yellow curve), [0.5, 4] *Hz* (Delta, red curve), [4, 8] *Hz* (Theta, pink curve), [8, 12] *Hz* (Alpha, green curve), and [12, 30] *Hz* (Beta, cyan curve). The vertical axis indicates true positive rates (TPRs), and the horizontal axis shows false positive rates (FPRs). The black line indicates the positive predictive value of 0.50 (PPV), and the region above this line shows the operating points with PPV ≥ 0.50. The right panel illustrates the zoomed-in version of the ROC curves around the optimal validation operating points across different frequency bands. The best SD detection performance corresponds to the Delta band, followed by Theta, near-DC, Alpha, and Beta band.

### D. Performance of WAVEFRONT in prediction of SD frequency

In this section, we explore the performance of our method in prediction of the number of SDs from the total minutes of detected SD events. This analysis is inspired by the recent work of Jewell *et al*. [10], where a linear regression was used for estimation of the number of SDs in 24-hour time windows. The frequency of occurrence of SDs could help clinicians make an informed decision on the type of medications and/or invasive procedures. High frequency/occurrence of SDs in continuous recordings of the brain is correlated with worsening brain injuries and poor outcomes in acute neurological conditions such as hemorrhage, ischemic stroke, and TBI [5, 10, 32, 60]. Therefore, measuring the frequency and duration of SDs is an important step toward personalized medicine.

The 74% detection rate of WAVEFRONT for SD events (see Section III-B for details) is promising, but is it sufficient for noninvasive estimation of the frequency of SDs? To evaluate WAVEFRONT’s performance for this purpose, we perform the following steps: (i) we extract overlapping time windows of 30-hour long, with a step size of 1 hour, across all of the 12 patients. Time windows with poor EEG quality are ignored, i.e., windows with less than 20 hours of reliable (not *masked out*, see Section II) EEG signals across ≥ 5 ipsilateral scalp electrodes. There are *N*_*w*_ =153 total time windows with *good* EEG quality (based on the definition provided above) in this dataset, (ii) for each time window, we apply WAVEFRONT to obtain the temporal detection output *T*_*out*_ (see Section II for details), where *T*_*out*_ = 1 indicates detected SD events at the corresponding time point, (iii) we prune and stitch together the detected intervals in each of the 30-hour windows. We remove isolated small detection intervals, which are less than 20min long and separated from other detection events with more than 4-hour temporal distance. After this pruning step, the remaining detected intervals are stitched together in a 4-hour sliding time window. This stitching process is at a very large temporal scale, in comparison to the original stitching process in the last step of WAVEFRONT (see Section II for details). The reason is that we are not looking for the single-SD detection performance in this analysis, but the performance of WAVEFRONT in prediction of SD frequency using the total duration of detections. Fig. 13 shows the total duration of detected events for each of the 153 time windows (blue dots), as a function of the number of annotated SD events. The number of SD events in these time windows ranges from 0 to 75, and the total detection duration in each time window lies in the range of 0 to 26.85 hours. In this figure, the increasing trend of detection duration as a function of the frequency of SDs is as expected. In addition, some piecewise flat parts around intervals of 22 to 37, and 54 to 75 SDs are observed in Fig. 13, which are resulting from underdetection of SD events in the time intervals, and (iv) finally, we measure the performance of WAVEFRONT in the estimation of the n<u>u</u>mber of SDs from the total duration of detected SD events in the 30-hour time intervals. A square root regression model of *a x* + *b* was used, where the number of annotated SDs in each window is the independent variable, and the total duration of detections is the observation. We choose a square root model due to the sublinear increasing trend of the total duration of detections for the larger numbers of SDs (e.g., ≥ 25 SDs). This sublinear increase rate of detection durations is caused by the underdetection of SDs, especially the highly clustered events, which underscores the limitation of WAVEFRONT in individual detection of SDs in these highly clustered events. We fit the square root model to the observations using a least square regression.

**Fig. 13.**
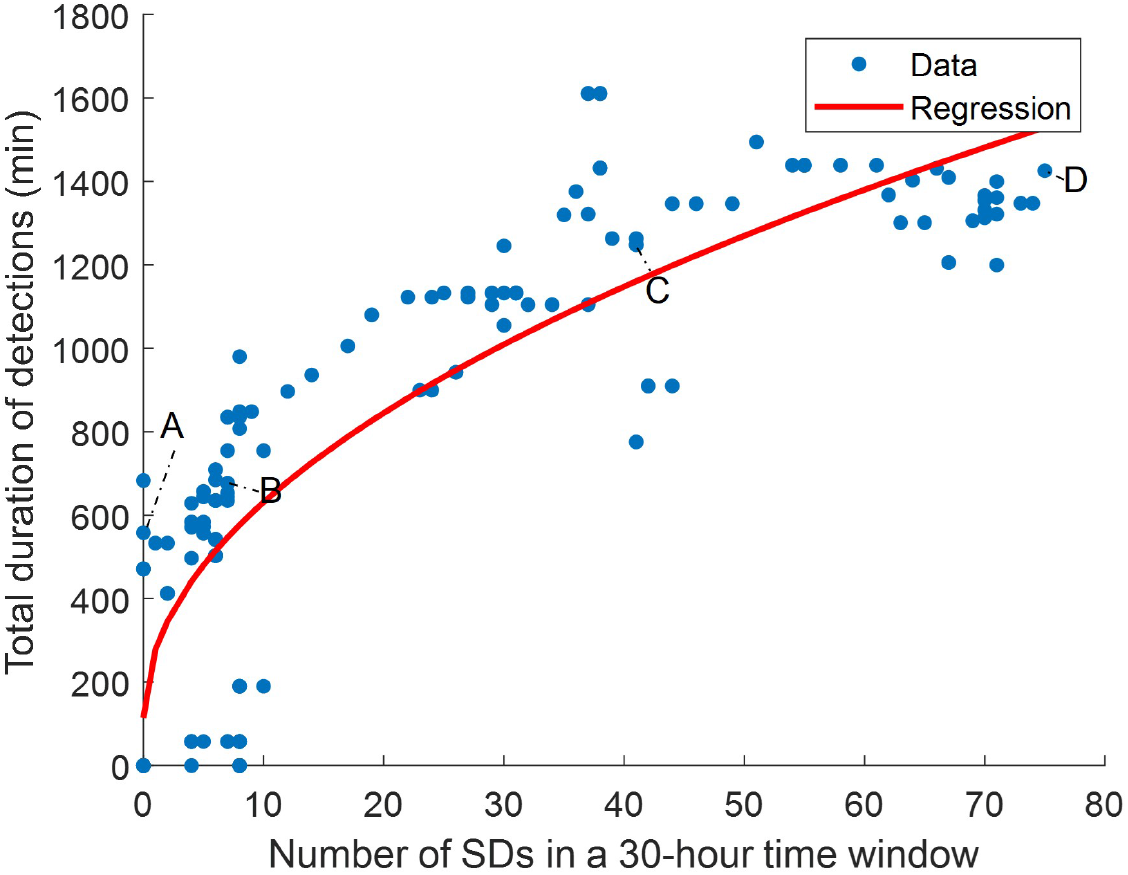
Performance of WAVEFRONT in prediction of SD frequency. Each blue dot shows the total duration of detected SD events using WAVEFRONT in 30-hour time windows, after pruning small and isolated detection events and stitching together the remaining detection events. The expected increasing trend of total detection duration as a function of the number of SD events is observed, with piecewise flat parts around 22-37, and 54-75 SDs which are the clustering detection side effects in these time intervals. A square root regression model (red curve) is fitted and used to quantify the prediction performance (R^2^ ≃ 0.71). Detected intervals in windows with small (point A and B), medium (point C), and large (Point D) number of SDs are shown in Fig 14.

Preliminary results, albeit with limited data, suggest that WAVEFRONT can reliably estimate the number of occurrences of SDs in long time intervals of 30-hour, with R^2^ ≃0.71 and 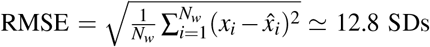 12.8 SDs, using a square root regression, as it is shown in Fig. 13. *x*_*i*_ is the number of annotated SDs in the i-th window, and 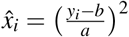 is the estimated number of SDs through the regression analysis and based on the total duration of detections in each time window (i.e., *y*_*i*_). Based on the results, WAVEFRONT can successfully discriminate between long windows of recordings with a large number of SDs (e.g., *>* 40) and a small number of SDs (e.g., *<* 20), and estimate the frequency of SDs with RMSE of less than 13 SDs. Such an analysis could also be used to assgin the patient an “SD-score”. More accurate estimation of number of SD events requires further improvements in the algorithm. Fig. 14 shows the detected intervals (marked with red strips on the bottom), along with the ground truth annotated SDs (dashed vertical lines) in some sample time windows with small, medium, and large number of SDs. Fig. 14a corresponds to point A in Fig. 13 and shows the detected false alarms in a patient without any SDs (patient 7), with a total detection duration of 558 min. The quality of the EEG recording in this time window is poor with large portions of the signals of 4 of the electrodes missing, which may explain the large number of false alarm detections. Fig. 14b corresponds to point B and shows a 30-hour time window with 7 SD events, where the WAVEFRONT algorithm successfully detected the isolated event, as well as the clustered scCSD events (see Section II for the definitions of SD annotations). Fig. 14c and d show time windows with larger number of SDs, in two patients with right (patient 3) and left (patient 12) DHC. These windows correspond to points C and D in Fig.13), with 41 and 75 SDs respectively, with highly clustered detection intervals. There are some missed SDs in between the two detection intervals in Fig. 14d, which maybe due to the removed outlier artifact across all of the EEG electrodes.

**Fig. 14.**
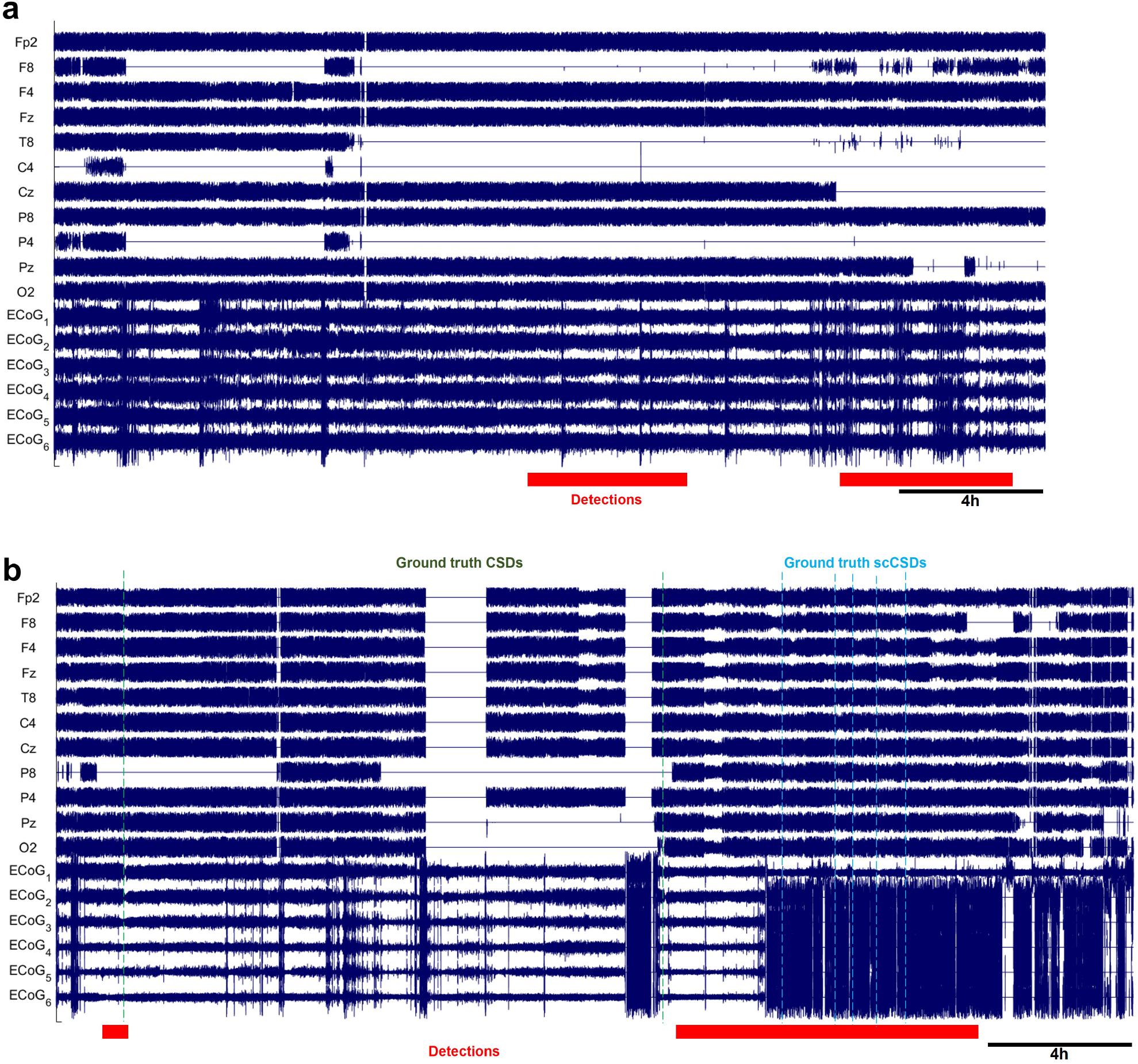
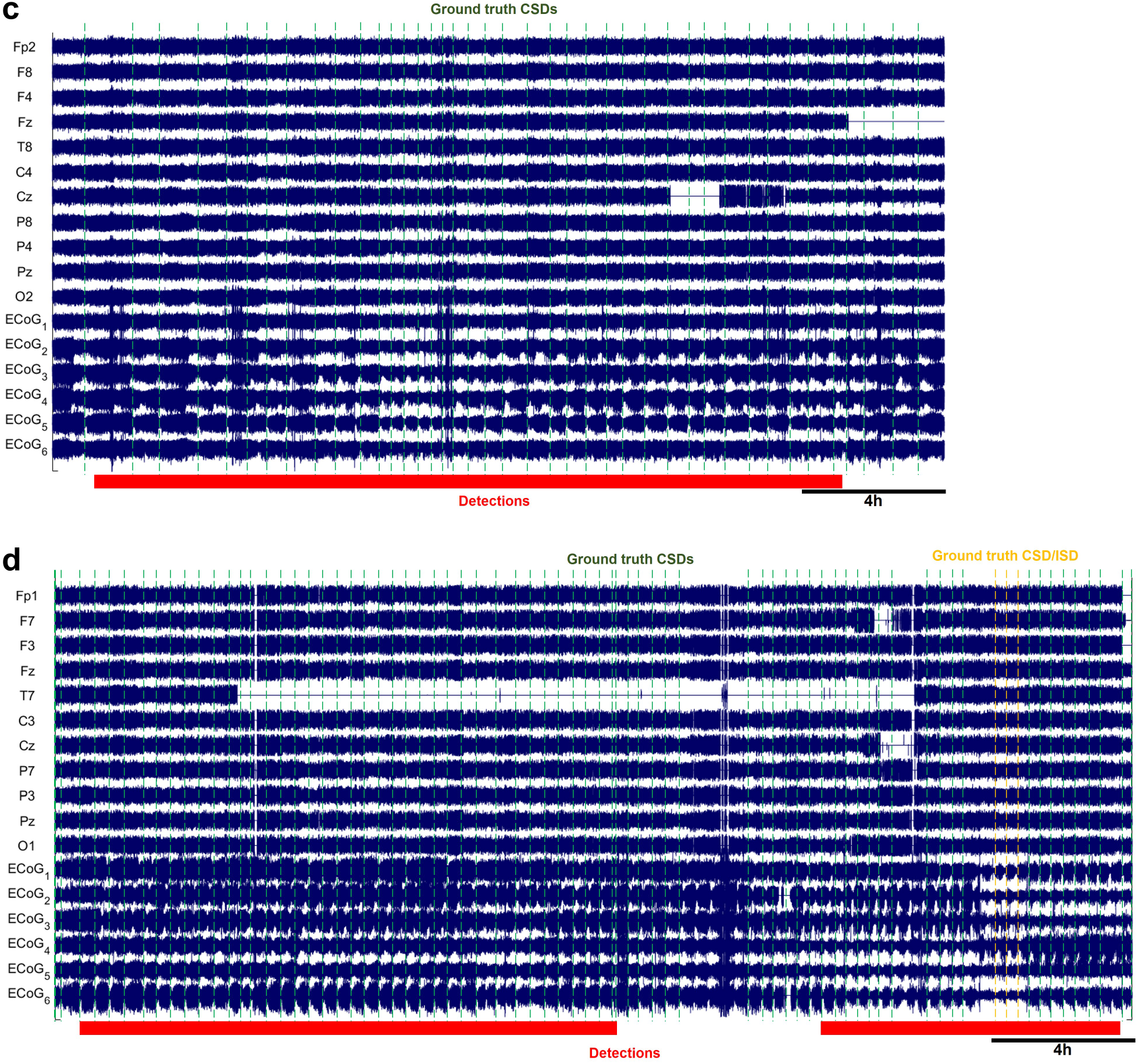
Detection intervals (marked with red strips on the bottom), along with the ground truth annotated SDs (dashed vertical lines) in long time windows, where the preprocessed ipsilateral EEG signals, along with the ECoG signals are shown. EEG electrodes are ordered using the transverse montage (see Fig. 10a). Signals are normalized by their standard deviations for the illustration purpose: a) A 26.6-hour time window, corresponding to point A in Fig. 13, with no SD event in patient 7. However, there are large false alarm detection intervals with total duration of 558min, which may be explained by the poor quality of the EEG recording in this time window, b) A 30-hour time window with 7 SDs (2 CSDs and 7 scCSDs), which corresponds to point B in the regression figure, and recorded from patient 6. WAVEFRONT successfully detected the the isolated CSD event, as well as the clustered SD events toward the end of the window, with the total detection duration of 676min. (Cont.). c) A 24.84-hour time window with highly clustered 41 SDs (point C in Fig. 13) in patient 3, and d) A 30-hour time window with the largest number of 75 SD events (point D in Fig. 13) in patient 12. WAVEFRONT detects long intervals in (c) and (d), with the total duration of 1248min and 1425min.

## IV. DISCUSSIONS AND CONCLUSIONS

In this study, we explored the feasibility and quantified the performance of automated noninvasive SD detection using continuous scalp EEG recordings from 12 severe TBI patients. These patients underwent DHCs, and experienced 700 total SD propagation events over days (95±42.2 hours) of simultaneous EEG and ECoG recordings at ICUs. Intracranial signals were used for SD event temporal annotation. Our previously proposed WAVEFRONT algorithm [28], with appropriate modifications and improvements, achieved a reliable SD detection performance of 74% average cross-validation TPR (∼13k of the ∼17k total SD windows across the validation sets are detected), with less than 1.5% average cross-validation FPR (less than 7k false alarms among the total 450k non-SD intervals) using Delta band scalp EEG signals. For the two patients without any annotated SD, the average false alarm rate was 1.7%, similar to the overal average of 1.5% cross-validation FPR. To understand the clinical implications of this, we evaluated the performance of WAVEFRONT in prediction of the number of SDs in long time intervals of 30-hour, using a square root regression. Preliminary results, albeit with limited data, suggest that WAVEFRONT achieves a promising performance (regression with R^2^ ≃ 0.71) in the estimation of SD frequencies, despite a substantial number of false alarms. The SD detection performance in Delta band was better than that in Theta (73% TPR, 2.0% FPR), near-DC (70% TPR, 2.1% FPR), Alpha (65% TPR, 2.0% FPR), and Beta (59% TPR, 1.6% FPR) bands. This decreasing trend of SD detection performance is consistent with existing understanding and literature, as the depth of SD depressions (i.e., percentage of maximum power reduction from the baseline power) reduces in higher frequency bands, with the largest reported depression depth, and highest baseline level in the Delta band EEG [22]. However, although SPCs are the prominent signatures of SDs in DC or near-DC ECoG signals [8], WAVEFRONT’s performance is worse when using near-DC EEG, in comparison to the higher frequency components in Delta band. This may be due to the inherent limitation of EEG in capturing the slow DC shifts [22], or the limitation of WAVEFRONT in the extraction of SPCs. The estimated average propagation speed of the detected SD events in the EEG Delta band using WAVEFRONT is 3.35±0.05 mm/min.

The WAVEFRONT algorithm achieved a reliable automated detection performance using scalp EEG. Preliminary evidence in SD frequency analysis suggests that WAVEFRONT achieves a promising performance in the prediction of SD frequency in long time intervals. Increasing evidence shows that SDs are the reliable predictor of outcomes in TBI patients [5, 10, 19, 33]. WAVEFRONT can potentially be used for prognostication of worsening brain injuries by providing a measure of SD frequency. SD detection on patients with DHC is a clinically relevant problem. DHC is widely used for the management of severe TBI patients [35–37]. Around 60% of these patients experience SDs, mostly with a high frequency of occurrence [10, 32], which significantly increases the chance of worsening brain injuries [5, 10, 19]. Every year in the United States, more than 1.2 million TBI patients experience worsening brain injuries or death [21], who form a large target population for continuous monitoring of SDs at ICUs. Scalp EEG in DHC patients provides wider spatial coverage of the brain than a locally placed intracranial strip of electrodes on the cortex, has a higher spatial resolution in comparison to the EEG recordings from patients with intact skull [35], and is less risky than intracranial electrode placement. Therefore, noninvasive detection and monitoring of SDs in severe TBI patients with DHC can help improve outcomes. The extension of the results in this paper to patients with intact skulls requires further study.

However, there are limitations associated with the SD ground truth and the dataset in this study: (i) due to limited spatial coverage of the strip of ECoG electrodes, the temporal annotations may not reflect the actual temporal onset of each SD event and some of the SDs may be even missed in the “ground truth” here, as some of the waves may have started to propagate from an origin far from the intracranial strip. However, ipsilateral EEG electrodes provide a full spatial coverage of the DHC hemisphere. This ground truth limitation makes it infeasible to quantify the performance of WAVEFRONT in determining whether SDs are present or not in each window. This may explain the limitation of WAVEFRONT in discrimination between windows with and without SDs, as it is shown in Fig. 13. Another contributing factor to this poor discrimination performance might be the inherent limitation of WAVEFRONT algorithm. This requires further study with higher spatial coverage of subdural SD recordings, (ii) the depression width, temporal duration, and propagation speed of SD waves are unknown. This may have resulted in slight over or underestimation of the actual performance of WAVEFRONT in the detection of SD events. A ground truth with higher spatial and temporal coverage and accuracy is warranted to further assess the performance of WAVEFRONT, (iii) due to the small number of patients in this study, overfitting to the available SD events is inevitable (see Section III for more details on this issue). We expect WAVEFRONT to achieve a better average validation performance by using a larger dataset of TBI patients with multiple SD events across different varieties of propagation patterns (single-gyrus, semi-planar, ring-shape, etc.), different ranges of propagation speeds, and in different brain regions. In addition, we would be able to provide statistical guarantees for the detection and discrimination results using a larger dataset, and (iv) low-density EEG with a small number of electrodes was used in this dataset, which limits the performance of WAVEFRONT. Based on our reported simulation results in [28], WAVEFRONT can detect narrow SD wavefronts, even single-gyrus SD propagations, using a sufficiently high density of EEG electrodes on the scalp. Thus, higher density EEG might be needed for milder TBIs with narrow SD wavefronts.

In addition to the limitations of the dataset and ground truth, WAVEFRONT has inherent limitations: (i) this algorithm suffers from underdetection of SD events since clustered SDs (more than two SDs in a time interval of 3 hours or less [10, 19]) in EEG cannot be detected individually using our current approach, (ii) although the average false alarm rate of ∼1.5% may seem small, this number corresponds to the ∼33% of the total detected events, i.e., out of the total detected events, around one-third of them are false alarms. This can be a serious limitation for diagnostic and monitoring goals, including the cases when the risk and side effects of interventions and treatments are high and a much lower false alarm rate is required, and (iii) finding the right set of stitching parameters in the SD frequency analysis is heuristic and there is room for improvements. This is a future direction for this work. Again, increase in the size of the dataset can help reduce FPR further.

This work is the first attempt to explore the feasibility and quantify the reliability of noninvasive SD detection in severe TBI patients using an automated algorithm, which can potentially be used for prognostication of worsening brain injuries, and paves the way toward personalized medicine.

## ACKNOWLEDGEMENTS

This work was supported, in part, by grants from the National Science Foundation (NSF), the Chuck Noll Foundation for Brain Injury Research, the Center for Machine Learning and Health at CMU, under the Pittsburgh Health Data Alliance, the Neil and Jo Bushnell Fellowship in Engineering, the Hsu Chang Memorial Fellowship, the CMU Swartz Center for Entrepreneurship Innovation Commercialization Fellows program, and by the Office of the Assistant Secretary of Defense for Health Affairs, through the Defense Medical Research and Development Program under Award No. W81XWH-16-2-0020. Dr. Elmer’s research time was supported by the National Institutes of Health (NIH) through grant 5K23NS097629. Opinions, interpretations, conclusions, and recommendations are those of the authors and are not necessarily endorsed by the Department of Defense. We thank Maysamreza Chamanzar, Shilpa George, Neil Mehta, David Okonkwo, and Praveen Venkatesh for helpful discussions.

## Footnotes

1 Centers for Disease Control and Prevention (CDC) National Center for Health Statistics.

2 In [10], the overall sensitivity was calculated by comparing the number of detected SDs against the number of ground truth SDs in 24-hour non-overlapping time windows across patients. They used a linear regression for this comparison with a slope of ∼79%. The reported false positive rate (FPR) is a median value of the calculated FPR for each of the 24-hour time windows, where the negative events are defined as 20min periods without any ground truth SDs.

3 ClinicalTrials.gov Protocol ID: 08-96-12-01

4 Prof. Jed A. Hartings at Department of Neurosurgery, University of Cincinnati

## REFERENCES

[1] G. G. Somjen. Ions in the brain: normal function, seizures, and stroke. Oxford University Press, 2004.

[2] B.J. Zandt, B. ten Haken, M. J. van Putten, and M. A. Dahlem. How does spreading depression spread? physiology and modeling. Reviews in the Neurosciences, 26(2):183–198, 2015.

[3] J. P. Dreier et al. Delayed ischaemic neurological deficits after subarachnoid haemorrhage are associated with clusters of spreading depolarizations. Brain, 129(12):3224–3237, 2006.

[4] J. P. Dreier. The role of spreading depression, spreading depolarization and spreading ischemia in neurological disease. Nature Medicine, 17(4):439–447, 2011.

[5] M. Lauritzen et al. Clinical relevance of cortical spreading depression in neurological disorders: migraine, malignant stroke, subarachnoid and intracranial hemorrhage, and traumatic brain injury. J. Cereb. Blood Flow Metab, 31(1):17–35, 2011.

[6] J. P. Dreier et al. Cortical spreading ischaemia is a novel process involved in ischaemic damage in patients with aneurysmal subarachnoid haemorrhage. Brain, 132(7):1866–1881, 2009.

[7] J. A. Hartings et al. Spreading depolarizations and late secondary insults after traumatic brain injury. Journal of neurotrauma, 26(11):1857–1866, 2009.

[8] J. P. Dreier et al. Recording, analysis, and interpretation of spreading depolarizations in neurointensive care: Review and recommendations of the cosbid research group. Journal of Cerebral Blood Flow & Metabolism, 37(5):1595–1625, 2017.

[9] S. Sivakumar et al. Cortical spreading depolarizations and clinically measured scalp eeg activity after aneurysmal subarachnoid hemorrhage and traumatic brain injury. Neurocritical Care, pages 1–11, 2022.

[10] S. Jewell et al. Development and evaluation of a method for automated detection of spreading depolarizations in the injured human brain. Neurocritical care, 35(2):160–175, 2021.

[11] R Sánchez-Porras et al. The role of spreading depolarization in subarachnoid hemorrhage. European Journal of Neurology, 20(8):1121–1127, 2013.

[12] J. Woitzik et al. Delayed cerebral ischemia and spreading depolarization in absence of angiographic vasospasm after subarachnoid hemorrhage. Journal of Cerebral Blood Flow & Metabolism, 32(2):203–212, 2012.

[13] K. Sugimoto and D. Y. Chung. Spreading depolarizations and subarachnoid hemorrhage. Neurotherapeutics, 17(2):497–510, 2020.

[14] H. Lantigua et al. Subarachnoid hemorrhage: who dies, and why? Critical care, 19(1):1–10, 2015.

[15] Y. Roos et al. Complications and outcome in patients with aneurysmal subarachnoid haemorrhage: a prospective hospital based cohort study in the netherlands. Journal of Neurology, Neurosurgery & Psychiatry, 68(3):337–341, 2000.

[16] D. Y. Chung, F. Oka, and C. Ayata. Spreading depolarizations: a therapeutic target against delayed cerebral ischemia after subarachnoid hemorrhage. Journal of clinical neurophysiology: official publication of the American Electroencephalographic Society, 33(3):196, 2016.

[17] B. Bosche et al. Recurrent spreading depolarizations after subarachnoid hemorrhage decreases oxygen availability in human cerebral cortex. Annals of neurology, 67(5):607–617, 2010.

[18] D. R. Kramer, T. Fujii, I. Ohiorhenuan, and C. Y. Liu. Cortical spreading depolarization: pathophysiology, implications, and future directions. Journal of Clinical Neuroscience, 24:22–27, 2016.

[19] D. Hertle et al. Effect of analgesics and sedatives on the occurrence of spreading depolarizations accompanying acute brain injury. Brain, 135(8):2390–2398, 2012.

[20] A. P. Carlson, M. Abbas, R. L. Alunday, F. Qeadan, and C. W. Shuttleworth. Spreading depolarization in acute brain injury inhibited by ketamine: a prospective, randomized, multiple crossover trial. Journal of neurosurgery, 130(5):1513–1519, 2018.

[21] Centers for Disease Control and Prevention (CDC), https://www.cdc.gov/traumaticbraininjury/moderate-severe/index.html.

[22] J. A. Hartings et al. Spreading depression in continuous electroencephalography of brain trauma. Annals of neurology, 76(5), 2014.

[23] C. Drenckhahn et al. Correlates of spreading depolarization in human scalp electroencephalography. Brain, 135(3):853–868, 2012.

[24] Z. JR. Bastany et al. Association of cortical spreading depression and seizures in patients with medically intractable epilepsy. Clin. Neurophysiol., 131(12):2861–2874, 2020.

[25] J. Hofmeijer et al. Detecting cortical spreading depolarization with full band scalp electroencephalography: an illusion? Frontiers in neurology, 9:17, 2018.

[26] J. A. Hartings, Laura B Ngwenya, Tomas Watanabe, and Brandon Foreman. Commentary: detecting cortical spreading depolarization with full band scalp electroencephalography: an illusion? Frontiers in Systems Neuroscience, 12:19, 2018.

[27] A. Chamanzar et al. Automated, scalable and generalizable deep learning for tracking cortical spreading depression using EEG. In 2021 10th International IEEE/EMBS Conference on Neural Engineering (NER), pages 416–419. IEEE, 2021.

[28] A. Chamanzar et al. An algorithm for automated, noninvasive detection of cortical spreading depolarizations based on EEG simulations. IEEE. Trans. Biomed. Eng., 66(4):1115–1126, 2018.

[29] S. J. Hund et al. Numerical simulation of concussive-generated cortical spreading depolarization to optimize dc-eeg electrode spacing for noninvasive visual detection. Neurocritical Care, pages 1–16, 2022.

[30] A. Chamanzar and P. Grover. Silence localization. In 2019 9th International IEEE/EMBS Conference on Neural Engineering (NER), pages 1155–1158. IEEE, 2019.

[31] A. Chamanzar, M. Behrmann, and P. Grover. Neural silences can be localized rapidly using noninvasive scalp EEG. Communications biology, 4(1):1–17, 2021.

[32] J. A. Hartings et al. Prognostic value of spreading depolarizations in patients with severe traumatic brain injury. JAMA neurology, 77(4):489–499, 2020.

[33] J. A. Hartings et al. Surgical management of traumatic brain injury: a comparative-effectiveness study of 2 centers. Journal of Neurosurgery, 120(2):434–446, 2014.

[34] G. WJ. Hawryluk et al. Guidelines for the management of severe traumatic brain injury: 2020 update of the decompressive craniectomy recommendations. Neurosurgery, 87(3):427–434, 2020.

[35] B. Voytek et al. Hemicraniectomy: a new model for human electrophysiology with high spatio-temporal resolution. Journal of cognitive neuroscience, 22(11):2491–2502, 2010.

[36] J. Hempenstall, A. Sadek, and C. A. Eynon. Decompressive craniectomy in acute brain injury–lifting the lid on neurosurgical practice. Journal of the Intensive Care Society, 13(3):221–226, 2012.

[37] G. A. Grindlinger, D. H. Skavdahl, R. D. Ecker, and M. R. Sanborn. Decompressive craniectomy for severe traumatic brain injury: clinical study, literature review and meta-analysis. Springerplus, 5(1):1–12, 2016.

[38] C. Drenckhahn et al. Complications in aneurysmal subarachnoid hemorrhage patients with and without subdural electrode strip for electrocorticography. Journal of Clinical Neurophysiology, 33(3):250–259, 2016.

[39] J. Woitzik et al. Propagation of cortical spreading depolarization in the human cortex after malignant stroke. Neurology, 80(12):1095–1102, 2013.

[40] E. Santos, R. Sánchez-Porras, O. W. Sakowitz, J. P Dreier, and M. A. Dahlem. Heterogeneous propagation of spreading depolarizations in the lissencephalic and gyrencephalic brain. Journal of Cerebral Blood Flow & Metabolism, 37(7):2639–2643, 2017.

[41] D. Kaufmann et al. Heterogeneous incidence and propagation of spreading depolarizations. Journal of Cerebral Blood Flow & Metabolism, 37(5):1748–1762, 2017.

[42] E. Santos et al. Radial, spiral and reverberating waves of spreading depolarization occur in the gyrencephalic brain. Neuroimage, 99:244–255, 2014.

[43] B. KP. Horn and B. G. Schunck. Determining optical flow. Artificial intelligence, 17(1-3):185–203, 1981.

[44] B. D. Lucas et al. An iterative image registration technique with an application to stereo vision. 1981.

[45] J. A. Hartings et al. The continuum of spreading depolarizations in acute cortical lesion development: examining Leao’s legacy. J. Cereb. Blood Flow Metab, 37(5):1571–1594, 2017.

[46] A. Delorme and S. Makeig. EEGLAB: an open source toolbox for analysis of single-trial EEG dynamics including independent component analysis. Journal of neuroscience methods, 134(1):9–21, 2004.

[47] J. W. Tukey. Exploratory data analysis. Reading, Mass. : Addison-Wesley Pub. Co., 1977.

[48] Outlier, from Wikipedia, https://en.wikipedia.org/wiki/Outlier.

[49] MATLAB and Signal Processing Toolbox Release 2015b, The MathWorks, Inc., Natick, Massachusetts, United States.

[50] D. J. McFarland et al. Spatial filter selection for EEG-based communication. Electroencephalography and clinical Neurophysiology, 103(3):386–394, 1997.

[51] P. Zhou. Numerical analysis of electromagnetic fields. Springer Science & Business Media, 2012.

[52] R. Keys. Cubic convolution interpolation for digital image processing. IEEE transactions on acoustics, speech, and signal processing, 29(6):1153–1160, 1981.

[53] MATLAB and Image Processing Toolbox Release 2018b, The MathWorks, Inc., Natick, Massachusetts, United States.

[54] D. L. Olson and D. Delen. Advanced data mining techniques. Springer Science & Business Media, 2008.

[55] B. Efron. Bootstrap methods: another look at the jackknife. In Breakthroughs in statistics, pages 569–593. Springer, 1992.

[56] B. Efron. Better bootstrap confidence intervals. Journal of the American statistical Association, 82(397):171–185, 1987.

[57] A. W. Moore. Cross-validation for detecting and preventing overfitting. School of Computer Science Carneigie Mellon University, 2001.

[58] R. Kohavi et al. Bias plus variance decomposition for zero-one loss functions. In ICML, volume 96, pages 275–83, 1996.

[59] T. Fawcett. ROC graphs: Notes and practical considerations for researchers. Machine learning, 31(1):1–38, 2004.

[60] J. A. Hartings et al. Spreading depolarisations and outcome after traumatic brain injury: a prospective observational study. The Lancet Neurology, 10(12):1058–1064, 2011.

